# A plastidial retrograde-signal potentiates biosynthesis of systemic stress response activators

**DOI:** 10.1101/2021.09.21.461262

**Authors:** Liping Zeng, Jin-Zheng Wang, Xiang He, Haiyan Ke, Mark Lemos, William M. Gray, Katayoon Dehesh

**Affiliations:** Institute for Integrative Genome Biology and Department of Botany and Plant Sciences, University of California, Riverside, CA 92521; Laboratory of Allergy and Inflammation, Chengdu third people’s hospital branch of National Clinical Research Center for Respiratory Disease, Chengdu 610031, China; Department of Plant and Microbial Biology, University of Minnesota, St. Paul, MN 55108

**Keywords:** MAPK3/6, MEcPP, N-hydroxy-pipecolic acid, pipecolic acid, plastidial retrograde-signal, PP2C.D1

## Abstract

- Plants employ an array of intricate and hierarchical signaling cascades to perceive and transduce informational cues to synchronize and tailor adaptive responses. Systemic stress response (SSR) is a recognized complex signaling and response network quintessential to plant’s local and distal responses to environmental triggers, however, the identity of the initiating signals has remained fragmented.
- Here, we show that both biotic (aphids and viral pathogens) and abiotic (high-light and wounding) stresses induce accumulation of the plastidial-retrograde-signaling metabolite, methylerythritol cyclodiphosphate (MEcPP), leading to reduction of the phytohormone, auxin, and the subsequent decreased expression of the phosphatase, *PP2C.D1*.
- This enables phosphorylation of mitogen-activated protein kinases (MAPK3/6), and the consequential induction of the downstream events ultimately resulting in biosynthesis of the two SSR priming metabolites, pipecolic- and N-hydroxy-pipecolic acid.
- This work identifies plastids as the initiation site, and the plastidial retrograde-signal, MEcPP as the initiator of a multi-component signaling cascade potentiating the biosynthesis of SSR activators, in response to biotic and abiotic triggers.

## Introduction

Dynamic organization of strata of intertwined signaling circuitries is fundamental to the integrity of cellular homeostasis in response to informational cues. Stress responses are induced via intricate and highly organized tiers of signaling cascades where the deactivation/activation of one component potentiates interaction and function of another. Uncovering the nature, the organization, and the operational mode of these tiered communication networks is one of the prime challenges of biology.

A well-recognized key mechanism in the transduction of intracellular signals in eukaryotic organisms is transmission of information via posttranslational protein modifications, most notably reversible protein phosphorylation carried out by protein kinases and protein phosphatases. Protein phosphatases are classified into three groups, among them type 2C protein phosphatases (PP2Cs), a structurally unique class of Mg^2+^-/Mn^2+^-dependent enzymes (Olsen *et al*., 2006; Moorhead *et al*., 2007; Fuchs *et al*., 2013). The Arabidopsis genome encodes eighty PP2Cs, nine of which belong to the D-subclade (Fuchs *et al*., 2013). Initial computational analyses of PP2C.D proteins identified a putative bipartite nuclear localization signal in all nine family members together with a potential transmembrane spanning region in PP2C.D1, D3, D4, D6, D7, and D9 (Schweighofer *et al*., 2004). Subsequent studies using protein-GFP reporters noted exclusive presence of PP2C.D2, D5, and D6 on the plasma membrane, detected D1, D3, and D4 in the nuclear and cytosolic compartments, and D8 in mitochondria (Ren *et al*., 2018). In addition to the distinct localization patterns, phosphatases are implicated in different functions including regulation of apical hook development (Sentandreu *et al*., 2011; Spartz *et al*., 2014), auxin-induced cell expansion (Spartz *et al*., 2014; Ren *et al*., 2018; Wang, J *et al*., 2020), leaf senescence (Xiao *et al*., 2015), immune responses (Couto *et al*., 2016), and altered intracellular responses to exogenous and endogenous stimulus via their nuclear/cytosolic interactions with mitogen-activated protein kinases (MAPKs) (Schweighofer *et al*., 2007; Umbrasaite *et al*., 2010; Galletti *et al*., 2011; Fuchs *et al*., 2013).

In Arabidopsis MAPKs are divided into four (A-D) groups (Ichimura *et al*., 2002). Group A includes MAPK3, MAPK6, and their orthologs, activated by phosphorylation in response to biotic and abiotic stimuli and by developmental cues (Kiegerl *et al*., 2000; Zhang & Klessig, 2001; Ichimura *et al*., 2002; Seo *et al*., 2007). Specifically, a range of stressors such as producers of reactive oxygen species (ROS) trigger MAPKs activity leading to their transport to the nucleus where they reconfigure transcriptional landscape by phosphorylating transcription factors (Kovtun *et al*., 2000; Miles *et al*., 2005; Pitzschke & Hirt, 2009; Taj *et al*., 2010). Intriguingly, the activation of MAPK3/6 result in the induction of selected stress-response genes, and block the action of auxin, thus providing a link between oxidative stress and auxin signal transduction (Kovtun *et al*., 2000).

Auxin [indole-3-acetic acid (IAA)] is an indispensable morpho-regulatory hormone involved in integration of developmental and environmental signals into a complex regulatory network permitting optimal architectural modifications in response to the prevailing conditions (Gil *et al*.,2001; Cheong *et al*., 2002; Navarro *et al*., 2006; Spaepen *et al*., 2007; Kazan & Manners, 2009). As such, auxin homeostasis is key to refinement of plant responses to an array of environmental signals such as ROS (Laskowski *et al*., 2002; Zhong *et al*., 2010; Tognetti *et al*., 2012; Yu *et al*.,2013). Interestingly, the proposed connection between auxin and plastid-to-nucleus (retrograde) signaling implied primary function of plastidial retrograde signal in auxin-based signaling cascade (Glasser *et al*., 2014). Indeed, recently the methylerythritol phosphate (MEP)-pathway intermediate, methylerythritol cyclodiphosphate (MEcPP), was identified as the stress-specific retrograde signaling metabolite, modulating growth by reducing the abundance of auxin and its transporter PIN1 via dual transcriptional and posttranslational regulatory inputs in response to abiotic stresses (Jiang *et al*., 2018). The connection provided solid evidence for the primary role of plastids in establishing a balance between plant growth and stress responses in accordance to the prevailing conditions (Jiang *et al*., 2018; Jiang *et al*., 2019; Jiang *et al*., 2020). In addition, auxin is also a known key constituent of the phytohormone-based signaling network mediating the regulation of defense responses, as evident by the suppression of the majority of the auxin responsive genes after induction of systemic acquired resistance (SAR) (Wang *et al*., 2007; Verma *et al*., 2016).

Whole plant immunity, coined SAR, is the process of priming defense responses in leaves distal to the local infection (Hunt *et al*., 1996; Ryals *et al*., 1996). This process is central to a broad-spectrum immunity protecting plants from immediate and future biotic attacks (Pieterse *et al*.,2009; Leon-Reyes *et al*., 2010; Spoel & Dong, 2012). However, pathogens are not unique in their ability to elicit systemic signals, since abiotic stresses also induce the rapidly transmitted systemic signal(s) from local to distal leaves, a response known as systemic acquired acclimation (SAA), key to acclimatory responses and improved tolerance (Karpinski *et al*., 1999; Czarnocka *et al*., 2020; Zandalinas *et al*., 2020). It is of note that both SAR and SAA, the two seemingly independent systemic responses, are triggered by common stress signals such as ROS (Baxter *et al*., 2014). This is to be expected since the establishment of SAR is not independent of abiotic cues such as light (Zeier *et al*., 2004), suggestive of overlapping regulatory components between the two networks.

Long distance communication and signal amplification of SAR is triggered by a number of mobile metabolites including salicylic acid (SA), methyl salicylate (MeSA), a lipid-transfer protein designated defective in induced resistance, azelaic acid, glycerol-3-phosphate, pipecolic acid (Pip) and its derivative N-hydroxy-pipecolic acid (NHP) (Jung *et al*., 2009; Chanda *et al*.,2011; Navarova *et al*., 2012; Chen *et al*., 2018). Specifically, Pip and NHP are noted signaling molecules that control both SA-dependent and SA-independent SAR activation, and are indispensable for the establishment of nearly all the respective transcriptional responses (Bernsdorff *et al*., 2016). Pip is synthesized by the agd2-like defense response protein1 (ALD1) (Navarova *et al*., 2012; Ding *et al*., 2016; Hartmann *et al*., 2017). Subsequent N-hydroxylation of Pip by flavin-dependent monooxygenase 1 (FMO1) results in formation of NHP (Chen *et al*.,2018; Hartmann, M. *et al*., 2018). Ultimately, elevation of Pip and NHP levels enable the establishment of SAR associated priming responses (Navarova *et al*., 2012; Zeier, 2013; Ding *et al*., 2016; Chen *et al*., 2018). Interestingly however, a recent study shows that FMO1 also plays an important role in triggering of an SAA response, supporting the notion that SAA and SAR not only respond to the same triggers, but also share part or all steps of the same signaling pathway(s) (Baxter *et al*., 2014; Czarnocka *et al*., 2020).

Despite numerous reports on the establishment of systemic signaling, the identity and the complexity of the initiating signals potentiating this key defense/adaptive mechanism remains elusive. Here, the exploitation of genetically manipulated lines that either inducibly or constitutively (*ceh1* mutant) accumulate MEcPP, together with biotically and abiotically stressed plants accumulating MEcPP, aided us to uncover the organizational sequence and the mode of MEcPP-mediated action in potentiating a multi-component cascade responsible for the production metabolites that trigger a general systemic signaling responses (SSR). The sequence of these events commences by MEcPP-mediated reduction of auxin abundance and the consequential decreased expression of auxin response factors (*ARFs*), the transcriptional activators of *PP2C.D1*. The resulting reduction of *PP2C.D1* transcripts enables phosphorylation of MAPK3 and 6, and the consequential induction of events leading to biosynthesis of Pip and NHP, the two key activators of SSR triggered by abiotic (wounding and high light) and biotic (aphid and a vital pathogen) stresses.

Collectively our finding establishes MEcPP as an initiator of the SSR to defend plants against a myriad of environmental challenges.

## Materials and Methods

### Plant material and growth condition

*Arabdopsis thaliana* seedlings were grown in 16-h light/8-h dark cycles at ~22 °C. Two-week-old seedling were treated with DEX, MEcPP (100 μM), IAA (10 μm, 1hr), Luciferase (1mM), high light (800 μmol m^−2^sec^−1^) or wounded by forceps as described previously (Benn *et al*., 2016; Jiang *et al*., 2018). The *pp2c* mutant (SALK_099356) was obtained from TAIR. Seedlings were grown under 12h light photoperiod at 22-24 °C for aphids infestation.

### Phylogenetic analyses

Protein sequences of PP2C Clade-D family (Xue *et al*., 2008) were downloaded from Phytozome. The software MEGA (Kumar *et al*., 2018) and PhyML (Guindon *et al*., 2010) was performed to construct the phylogeny.

### Luciferase-Activity quantification

Luciferase activity signals were detected by a CCD camera (Wang *et al*., 2014). Quantification and statistical analyses of *RSRE:LUC* activity were performed (Benn *et al*., 2014).

### Metabolites extraction and analyses

MEcPP, IAA and Salicylic acid were analyzed as previously described (Jiang *et al*., 2019).

Pip measurements were performed using a Dionex Ultimate 3000 binary RSLC system coupled to Thermo Q-Exactive Focus mass spectrometer with a heated electro spray ionization source. Plant samples were separated using an Accucore-150-Amide-HILIC column (150 × 2.1 mm; particle size 2.6μM; Thermo Scientific 16726-152130) with a guard column containing the same column matrix (Thermo Scientific 852-00; 16726-012105). Gradient elution was carried out with acetonitrile (A) and 10 mM ammonium acetate pH 7.0 (B). The separation was conducted using the gradient profile (*t* (min), %A, %B): (−2, 90, 10), (0, 90, 10), (12, 30, 70), (15, 30, 70), (16, 90, 10), (22, 90, 10). The flow rate was kept at 280 μL/min and the injected volume was 2 μL. The column was kept at 35 °C. Mass spectra were acquired in positive mode under the following parameters: spray voltage, 4.50 KV; sheath gas flow rate 50, auxiliary gas flow rate 14, sweep gas flow rate 2, capillary temperature of 275 °C, S-lens RF level 100 and auxiliary gas heater temperature 275 °C. The initial 0.5 min of each run was sent to waste to avoid salt contamination of the MS. Compounds of interest were identified by accurate mass measurements (MS1), retention time and mass transitions monitoring. Pip was identified by using standard (Sigma, P45850). For relative quantitation, peak area for each compound (MS1; Thermo Trace Finder Software) was normalized to the initial fresh weight mass.

*N*-OH-Pip was measured using the same system as Pip measurements. Plant samples were separated by an Acquity UPLC HSS T3 column (1.8-μm, 150 × 2.1 mm) (Waters, part # 186003539). The mobile phases were A (ACN, 0.1% FA) and B (water, 0.1% FA), and the gradient was implemented at a flowrate of 0.2 mL/min (percentages indicate percent B): 0-1 min (99%), 1-8 min (99-50%), 8-10 min (50%), 10-10.5 min (50–99%), and 10.5-13 min (99%). The column was kept at 35 °C. The MS was run in positive ion mode with the following parameters: spray voltage, 4.50 KV; sheath gas flow rate 45, auxiliary gas flow rate 20, sweep gas flow rate 2, capillary temperature of 250 °C, S-lens RF level 50 and auxiliary gas heater temperature 250 °C. The initial 1 min of each run was sent to waste to avoid salt contamination of the MS. NHP was accurately identified by using the standard obtained from Professor Elizabeth Sattely’s lab at Stanford university, and by accurate mass measurements (MS1), retention time and mass transitions. For relative quantitation, peak area for each compound (MS1; Thermo Trace Finder Software) was normalized to the initial fresh weight mass.

### RNA-seq Analysis

Two-week-old *A. thaliana* seedlings were collected. RNA-seq libraries construction followed the BrAD-seq method (Townsley *et al*., 2015). Each genotype has six biological replicates. 75bases of single-end reads were sequenced. Tophat2 (Kim *et al*., 2013) was used to map reads to the genome of *A. thaliana*. DESeq2 (Love *et al*., 2014) was used to count and normalize mapped reads. Genes with 2-fold altered expression levels and p-value ≤0.05 were identified as differentially expressed genes. The GO term enrichment analyses were obtained by agriGOv2, and the heatmap was generated by the pheatmap (Kolde & Kolde, 2015) in *R* program (Team, 2013) (Table S2). RNA-seq data of SAR and Pip response genes were downloaded from published (Hartmann *et al*., 2017). List of genes with RSRE-containing promoters were obtained from published (Benn *et al*., 2016).

### Quantification of gene expression

RT-qPCR was performed as described previously (Walley *et al*., 2007). The control genes was AT4G26410. Table S3 listed primer sequences.

### Agro-infiltration-based transient assays in *Nicotiana benthamiana*

*N. benthamiana* transient assay was used to identify the protein-protein interaction between *PP2C* and *MAPK3*, and *MAPK6*. pENTR/D-TOPO (Invitrogen) and Gateway systems were used for constructing vectors. Vectors containing C/N-terminal luciferase fused with *PP2C, MAPK3* and *MAPK6* were introduced into *Agrobacterium* GV3101 and subsequently used for infiltration of *N. benthamiana* leaves, followed by luciferase activity signal detection using CCD camera (Wang *et al*., 2014).

### Co-Immunoprecipitation

Two-week-old seedlings were ground in liquid nitrogen and suspended in 2x extraction buffer (50 mM Tris-HCl at pH 7.5, 150 mM NaCl, 10% glycerol, 0.1% NP-40, protease inhibitor cocktail and phosphatase inhibitor) at 4 °C for 30 min. The protein suspensions were then centrifuged at 4,000*g* for 10 min and filtered out the precipitation using the 100 μm Nylon Mesh. The supernatant was incubated with GFP-Trap magnetic beads (for IP) and bmab-20 (negative control) (Chromotek), respectively, for 2 h at 4 °C. The beads were washed five times with the 2x extraction buffer. The immuno-precipitants were eluted with 2x SDS lysis buffer (50 mM Tris-HCl at pH 6.8, 2% SDS, 10% glycerol, 0.1% bromophenol blue, 1% 2-mercaptoethanol), boiled (100 °C, 10 min), separated on SDS-PAGE gel and subsequently transferred onto nitrocellulose membrane for probing with the corresponding antibodies.

### Protein extraction and immuno-blot analyses

Two-week-old seedlings were ground in liquid nitrogen and suspended in 2x SDS lysis buffer and boiled (100 °C, 10 min). Proteins were then separated on SDS-PAGE gel and transferred onto the nitrocellulose membrane. The monoclonal anti-PP2C (1:5000) was previously reported (Spartz *et al*., 2014). The phosphorylated MAPK3 and MAPK6 proteins were detected using polyclonal anti-phospho-p44/42 MAPK (Erk1/2, 1:1000; Cell Signaling Technology), and the detection of the total MAPK3 and 6 protein levels were by using polyclonal anti-MAPK3 (1:1000, Sigma) and anti-MAPK6 (1:1000, Sigma). The goat-anti-Rabbit (1:3000) HRP-conjugated secondary antibody was used.

### Aphid infestation

The potato aphids (*Macrosiphum euphorbiae*) isolate WU11 (Teixeira *et al*., 2018) were maintained on their adapted hosts for over 2.5 years in a growth chamber at 20°C with 16h light photoperiod. To infest new Arabidopsis seedlings, the colony was released to the growth chamber with the experimental plants to allow the infestation (Teixeira *et al*., 2018).

### Viral infection

Four-week-old seedlings were infected with *cucumber mosaic virus* (*CMV-m2b*) for 2 weeks.

### Accession Numbers and RNA-seq data

*PP2C.D1* (AT5G02760), *HDS* (AT5G60600), *MAPK3* (AT3G45640), *MAPK6* (AT2G43790), *SARD1* (AT1G73805), *CBP60g* (AT5G26920), *ALD1* (AT2G13810), *FMO1* (AT1G19250), *ARF7* (AT5G20730), *ARF19* (AT1G19220), *ARF2* (AT5G62000), *ARF3* (AT2G33860), *ARF10* (AT2G28350), *ARF11* (AT2G46530), *ARF18* (AT3G61830).

All RNA-seq data were submitted to NCBI SRA database (PRJNA596287).

## Results

### MEcPP-mediated transcriptional suppression of *PP2C.D1*

Comparative RNA-seq analyses of the high MEcPP-accumulating mutant, *ceh1*, versus wild-type plant revealed altered expression profile of the clade D phosphatases (Fig. **S1a-b**). Subsequent studies specifically identified *PP2C.D1*, also known as *APD7* (Arabidopsis PP2C clade D7) or *SSPP* (senescence-suppressed protein phosphatase) (Tovar-Mendez *et al*., 2014; Xiao *et al*.,2015), as the phosphatase with the most notably reduced transcript levels compared to the other clade members in the *ceh1* mutant. For simplicity throughout the paper, we will refer PP2C.D1 as PP2C. Indeed, qRT-PCR analyses confirmed markedly lower *PP2C* expression levels in *ceh1* compared with the wild-type (Fig. **1a**).

**Figure 1.**
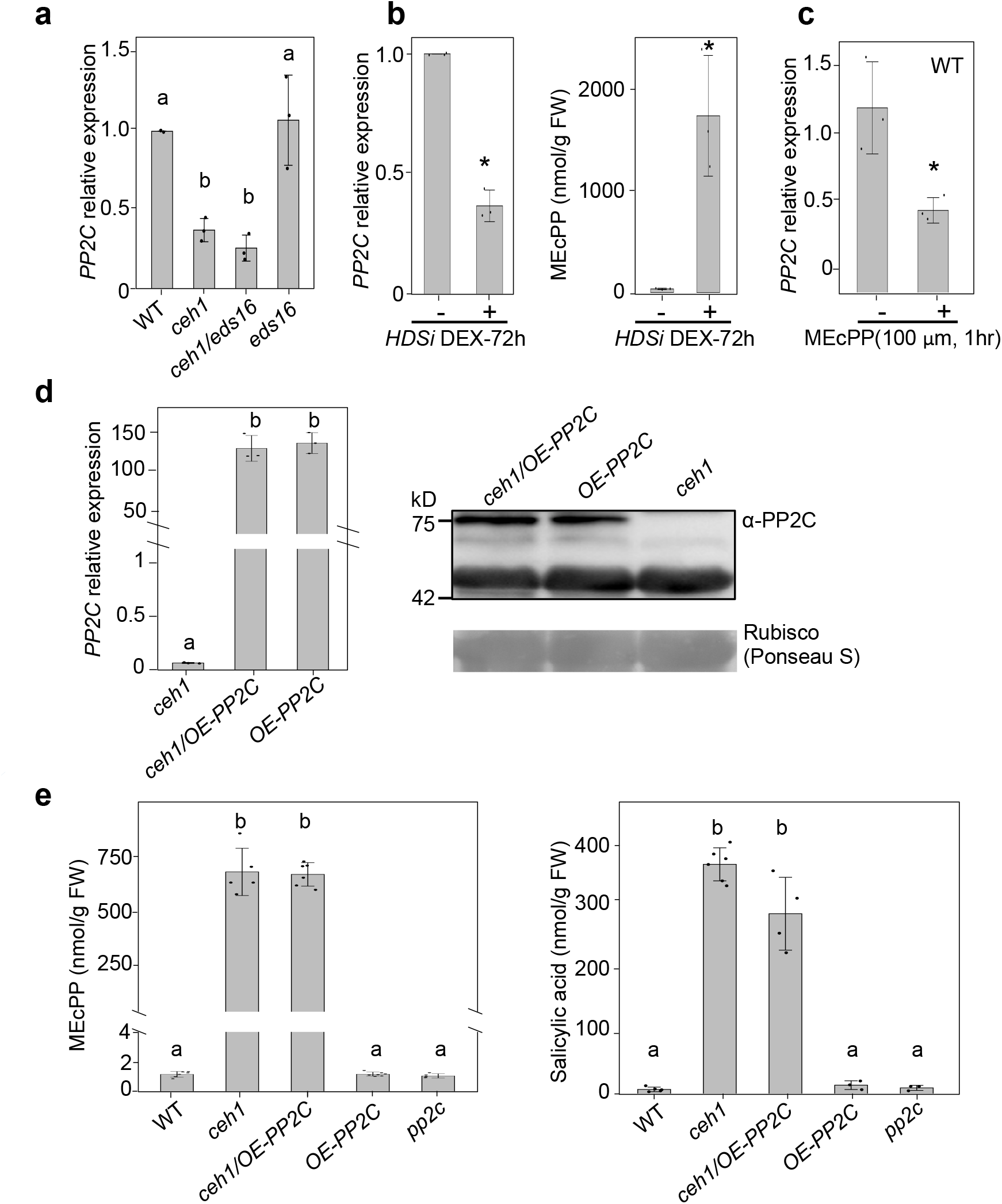
MEcPP-mediates transcriptional suppression of *PP2C*. (**a**) Suppression of *PP2C* expression is MEcPP-dependent and SA-independent. Total RNAs isolated from two-week-old seedlings of wild-type (WT), *ceh1/eds16, eds16* were subjected to qRT-PCR analyses. (**b**) Accumulation of MEcPP is inversely correlated to *PP2C* transcript levels. Relative expression of *PP2C* and MEcPP levels in DEX-inducible *HDSi* line 72 hours post mock- (-) and DEX-treatment (+). Analyses were performed on two-week-old seedlings. (**c**) Relative expression levels of *PP2C* in WT plants, 60 min post mock- (-) and MEcPP (100 μM)-treatment (+) confirms MEcPP-dependent transcriptional suppression of *PP2C*. (**d**) Reduced expression level of *PP2C* in *ceh1* is recovered in *PP2C* overexpressing *ceh1* (*ceh1/OE-PP2C*) and wild-type (*OE-PP2C*) lines. Immunoblot analyses using PP2C antibody display undetectable protein in *ceh1*, but detectably similar levels in PP2C overexpressing lines. The lower non-specific reacting band and Ponceau S staining show equal loading. (**e**) Analyses of MEcPP and SA levels in WT, *ceh1*, *ceh1/OE-PP2C*, *OE-PP2C* and *PP2C* mutant line (*pp2c*) show PP2C-independent accumulation of the metabolites. The *PP2C* mRNA levels was normalized to the levels of *At4g26410* (M3E9). All Data are mean ± SD of three biological and three technical replicates. Two-tailed Student’s *t* tests or ANOVA tests confirm MEcPP-mediated suppression of *PP2C*. Asterisks denotes significance. Lower case letters on top of histograms represent statistically significant differences (*P* ≤ 0.05).

Next, we analyzed the *PP2C* expression levels in salicylic acid deficient *eds16* and *ceh1/eds16* mutants to assess the potential regulatory input of the high SA present in *ceh1* mutant (Xiao *et al*.,2012) (Fig. **1a**). The results illustrate the SA-independent reduction of *PP2C* transcript levels in the high MEcPP-accumulating *ceh1* mutant backgrounds.

To investigate the MEcPP-mediated reduction of *PP2C* transcript levels, we exploited a dexamethasone (DEX)-inducible MEcPP accumulating line (*HDSi*), previously shown to accumulate similar MEcPP levels as that found in the *ceh1* mutant, at 72h post DEX-induction (Jiang *et al*., 2018; Jiang *et al*., 2019; Wang, JZ *et al*., 2020). The analyses of *PP2C* expression levels in mock- and DEX-treated plants (72h post induction) display an inverse correlation between DEX-inducible accumulation of MEcPP and expression levels of *PP2C* (Fig. **1b**).

To provide a direct evidence for MEcPP-mediated suppression of *PP2C* expression, we examined the relative transcript levels of the gene in mock- and exogenously MEcPP-treated wild-type plants (Fig. **1c**). Indeed, the reduced transcript levels of *PP2C* an hour post MEcPP application confirm specificity of MEcPP in mediating this suppression.

To assess whether MEcPP-mediated suppression of *PP2C* is via transcriptional and/or posttranscriptional modifications, we employed plants expressing *PP2C* under the control of the constitutive promoter, *35S:PP2C-GFP* (for simplicity herein designated *OE-PP2C*) and the introgressed line in the *ceh1* mutant background (*ceh1*/*OE-PP2C*) (Fig. **1d**). The similarly high *PP2C* transcripts in *OE-PP2C* and *ceh1*/*OE-PP2C* as compared to the notably reduced levels in the *ceh1* mutant background is a clear demonstration of the MEcPP-mediated transcriptional suppression of *PP2C*. Moreover, immunoblot analyses using PP2C specific antibody established the concordance between the protein and transcript levels, as evidenced by similarly high PP2C protein abundance in *OE-PP2C* and *ceh1*/*OE-PP2C* compared to undetectable protein levels in the *ceh1* mutant (Figs. **1d** **and S2**).

To examine a potential link between *PP2C* transcript levels and production of SA and/or MEcPP, we examined the abundance of these two metabolites in various genotypes (WT, *ceh1, ceh1/OE-PP2C, OE-PP2C*, and the *pp2c* mutant) (Fig. **1e**). The data explicitly confirm the PP2C-independent accumulation of MEcPP and SA.

Collectively, the finding establishes a SA-independent but MEcPP-dependent transcriptional suppression of *PP2C*, and excludes any PP2C regulatory input in accumulation of MEcPP and SA.

### MEcPP-mediated transcriptional regulation of *PP2C* is auxin-dependent

To dissect the regulatory components of *PP2C* transcriptional machinery, we examined and identified four auxin response *cis*-elements (AuxRE) (Ulmasov *et al*., 1995) in the *PP2C* promoter sequences (Fig. **2a**). The presence of these auxin-dependent regulatory elements together with reduced abundance of auxin and its transporter PIN1 (Jiang *et al*., 2018) via the MEcPP-mediated transcriptional and posttranslational regulatory input, led us to examine the *PP2C* transcript levels in mock- and auxin-treated *ceh1* and WT plants (Fig. **2b**). The elevated *PP2C* transcript levels in WT and the *ceh1* mutant an hour post IAA-treatment compared with untreated plants is an indicative of IAA-dependent transcriptional regulation of *PP2C* expression, corroborating the previous finding (Nemhauser *et al*., 2006; Ren *et al*., 2018). It is of note that lower levels of *PP2C* expression in IAA-treated *ceh1* relative to the corresponding WT is likely in part due to the impairment of auxin distribution in the *ceh1* mutant caused by reduced abundance of auxin transporter, PIN1 (Jiang *et al*., 2018). Alternatively and or additionally, the reduced expression of *PP2C* in IAA-treated *ceh1* relative to that of the WT plant could be attributed to decreased expression levels of auxin response factors (ARFs), a family of transcription factors responsible for the induction of AuxREs (Ulmasov *et al*., 1999; Guilfoyle & Hagen, 2001). To test this hypothesis, we examined expression levels of several family members of *ARFs* (Figs. **2c** and **S3**). Among the tested members, only *ARF7* and *19* displayed reduced transcript levels in *ceh1* backgrounds (*ceh1* and *ceh1/eds16*) compared to the corresponding controls (WT and *eds16*). This prompted us to examine the *PP2C* expression levels in mock- and auxin-treated single and double *arf7* and *19* mutants (Fig. **2d**). The partial induction of *PP2C* expression in auxin-treated single mutants as opposed to no induction in the double mutant line compared with the WT plant, establishes the key function of two auxin response factors, ARF7 and 19, in induction of *PP2C*.

**Figure 2.**
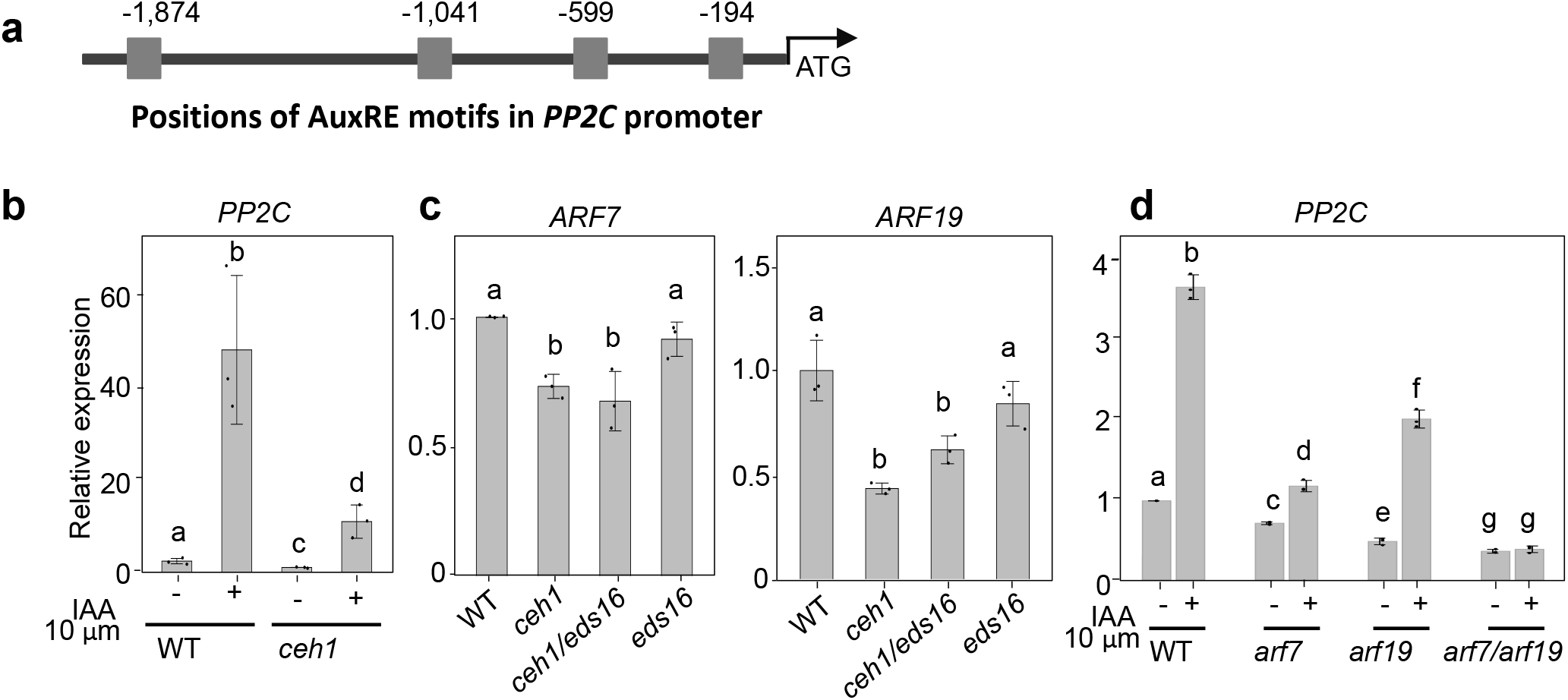
Auxin induces *PP2C* expression. (**a**) The schematic presentation of the *PP2C* promoter display the positions of auxin response elements (AuxRE). (**b**) Expression levels of *PP2C* an hour post mock- (-) and auxin (10 μM) - treatment (+) of *ceh1* and WT seedlings display enhanced expression in auxin-treated lines. (**c**) Reduced expression levels of auxin response factors (*ARF7* and *19*) in *ceh1* and *ceh1/eds16* compared to the levels in WT and *eds16* lines. (**d**) *PP2C* expression levels an hour post mock- (-) and auxin (10 μM)-treatment (+) of WT, single (*arf7*, and *arf19*) and double (*arf7/arf19*) mutant lines display indispensable function of ARF7 and 19 in auxin-induction of *PP2C* expression. Statistical analyses were performed by two-tailed Student’s *t* tests or ANOVA tests.

The data collectively delineate the molecular strata of MEcPP-mediated suppression of *PP2C* expression, commenced by reduced abundance of auxin, and the consequential decreased expression of *ARF7* and *19*, the *PP2C* transcriptional activators.

### *PP2C* suppresses the MEcPP-inducible RSRE-containing stress-response genes

To examine the consequences of altered *PP2C* transcript levels in high MEcPP containing plants, we analyzed the global expression profiles of *ceh1/OE-PP2C* and *HDSi/OE-PP2C* versus those corresponding to *ceh1* and *HDSi* backgrounds (Supplemental data sets **1-3**). The analyses revealed a notable presence (47-to-68%) of robustly suppressed genes in *ceh1/OE-PP2C* and *HDSi/OE-PP2C* backgrounds, respectively, that contain a general stress response (GSR) *cis*-element, coined *Rapid Stress Response Element* (*RSRE*) (Walley *et al*., 2007; Benn *et al*., 2014; Benn *et al*., 2016), in their promoters (Fig. **3a**, Table S**1**). To examine the validity of these analyses *in planta*, we employed *4xRSRE:Luciferase* line, used for functional readout of stress-induced rapid transcriptional responses (Walley *et al*., 2007; Benn *et al*., 2014; Bjornson *et al*.,2014), and introgressed it into the *ceh1* and *ceh1/OE-PP2C* backgrounds. Subsequent luciferase activity assays using homozygous introgressed lines showed markedly reduced luciferase activity in *ceh1/OE-PP2C/RSRE:LUC* line (herein designated as *ceh1/OE-PP2C*) compared with the previously established high and constitutive expression of the *RSRE* in *ceh1/RSRE:LUC* line (*ceh1*) (Benn *et al*., 2016) (Fig. **3b**). Moreover, additional bioinformatics analyses revealed a 27% increase in the number of induced stress-response genes containing RSRE in *pp2c* mutant compared to *OE-PP2C* line (Fig. 3**c**). Combined *in vivo* and in silico analyses support the involvement of PP2C in transcriptional regulation of RSRE-containing stress response genes.

**Figure 3.**
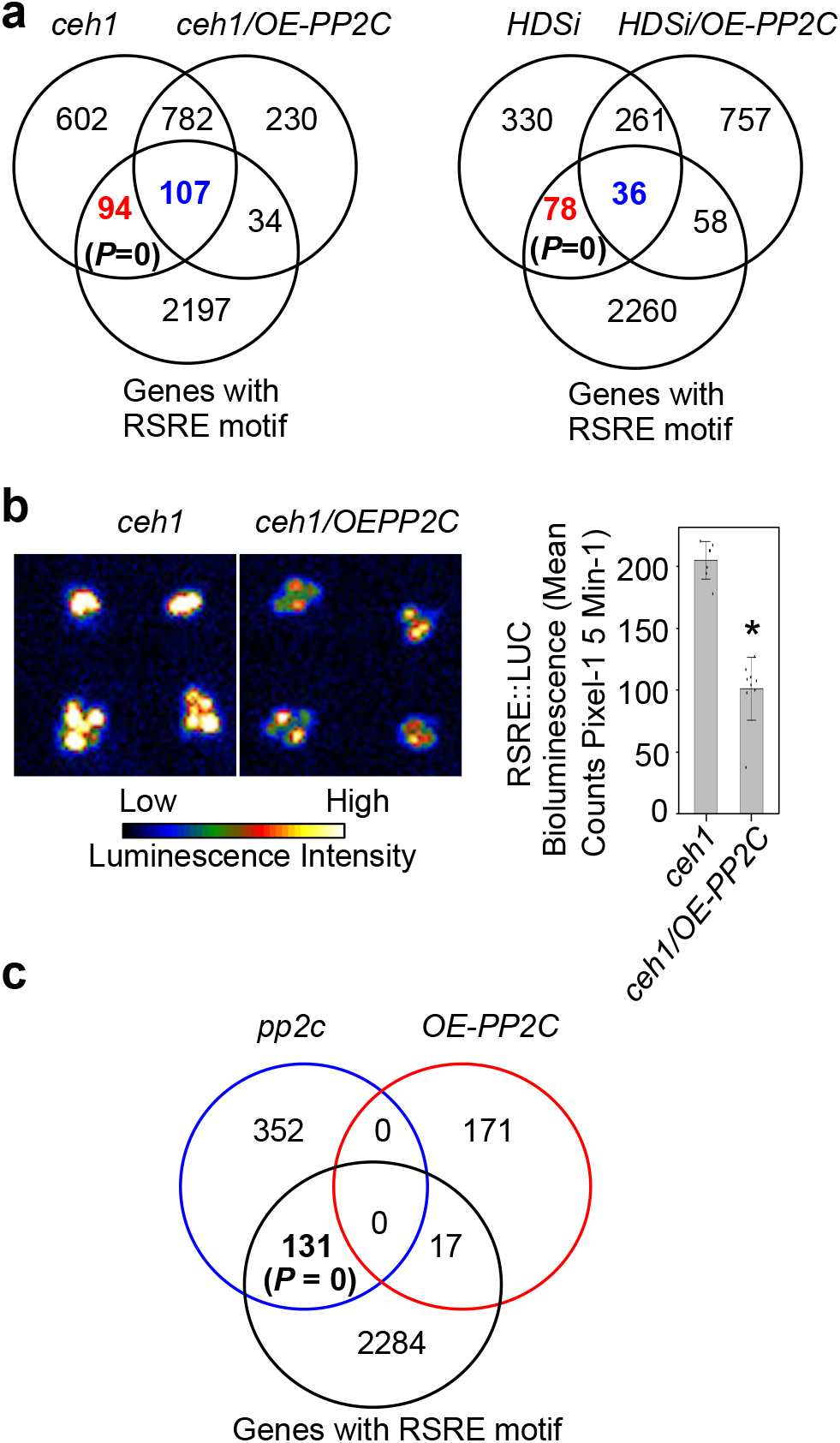
*PP2C* suppresses the MEcPP-inducible SAR and RSRE-containing genes. (**a**) Venn diagrams illustrate suppression of a significant number of MEcPP-inducible RSRE-motif containing genes in *PP2C* overexpressing (*ceh1/OE-PP2C and HDSi/OE-PP2C*) lines. (**b**) Representative images of *PP2C* overexpressing lines in the *ceh1* background (*ceh1/OE-PP2C*) show the notable reduction of RSRE:LUC activity compared to the *ceh1* mutant. The bar underneath displays the intensity of LUC activity and the histogram show quantification of LUC activity. Data are presented as means ± SEM. Asterisk notes statistically significant differences (*P* ≤ 0.05) by using two-tailed Student’s *t* test. (**c**) Venn diagrams illustrate enrichment of RSRE-motif containing genes whose expressions are significantly induced in the *pp2c* mutant but not in *OE-PP2C* plants under standard conditions.

Specifically, the Gene Ontology enrichment analyses of differentially expressed genes in *pp2c* mutant versus *OE*-*PP2C* lines revealed transcriptional profile that is partitioned into two distinct clusters of stress-response and growth-related genes (Fig. **S4** and Table S**2**). The inverse expression profiles of the two clusters support the notion of PP2C function in induction of growth-related genes, and suppression of stress-response genes (Fig. **S4**). Additional analyses established significant enrichment of induced plant-pathogen interaction pathway genes in *pp2c* mutant compared to the wild-type (Fig. **S5a**). This data support the recent report on reduction of *PP2C* transcript levels in response to flg22 and nlp20 treatment (Bjornson *et al*., 2021) (Fig. **S5b**).

Collectively, our findings uncover the PP2C-mediated transcriptional reconfiguration of RSRE containing genes, and further allude to the growth optimizing function of this phosphatase in concordance with its suggested role in promoting apical hook development in etiolated seedlings (Sentandreu *et al*., 2011; Spartz *et al*., 2014). Our experimental and bioinformatics data extend supports to the notion of PP2C function in both biotic and abiotic stress responses, and as a governing module balancing growth versus adaptive responses.

### PP2C suppresses transcription of Pip and NHP biosynthesis genes and their metabolites

Extended bioinformatics analyses unraveled a significant reduction in the number of SAR- (43-to-52%) and Pip-induced (44-to-40%) genes in *ceh1/OE-PP2C* and *HDSi/OE-PP2C* versus their corresponding backgrounds, respectively (Fig. **S6a-b** and Table **S1**). This together with the known functions of Pip and NHP in triggering SAR (Navarova *et al*., 2012; Hartmann, Michael *et al*., 2018), prompted us to test the potential involvement of MEcPP, SA, and PP2C in modulating the expression of genes involved in Pip and NHP biosynthesis. Specifically, we analyzed the relative expression levels of *SARD1, CBP60g, ALD1* and *FMO1* in WT, *ceh1, eds16*, *ceh1/eds16*, *HDSi*, *HDSi/eds16*, *PP2C* overexpressing WT (*OE-PP2C*), *ceh1* (*ceh1/OE-PP2C*) and *pp2c* lines (Fig. **4a-b**). Similar expression profiles of the aforementioned genes in the *ceh1* mutant and the DEX-induced *HDSi* line relative to the corresponding controls (WT and mock-treated *HDSi*) illustrate their MEcPP-dependent induction in constitutive and in inducible lines (Fig. **4b**). However, while MEcPP induces expression of all the genes (*SARD1*, *CBP60g*, *ALD1* and *FMO1*), SA differentially alters their expression profile, as evidenced by the reduced *SARD1* but induced *FMO1* expression levels in SA-deficient *ceh1/eds16* line compared to *ceh1*. Moreover, the SA-mediated reduction of *CBP60g* in the inducible line is hindered by constitutive production of MEcPP, whereas the *ALD1* transcript levels remain SA-independent (Fig. **4b**).

**Figure 4.**
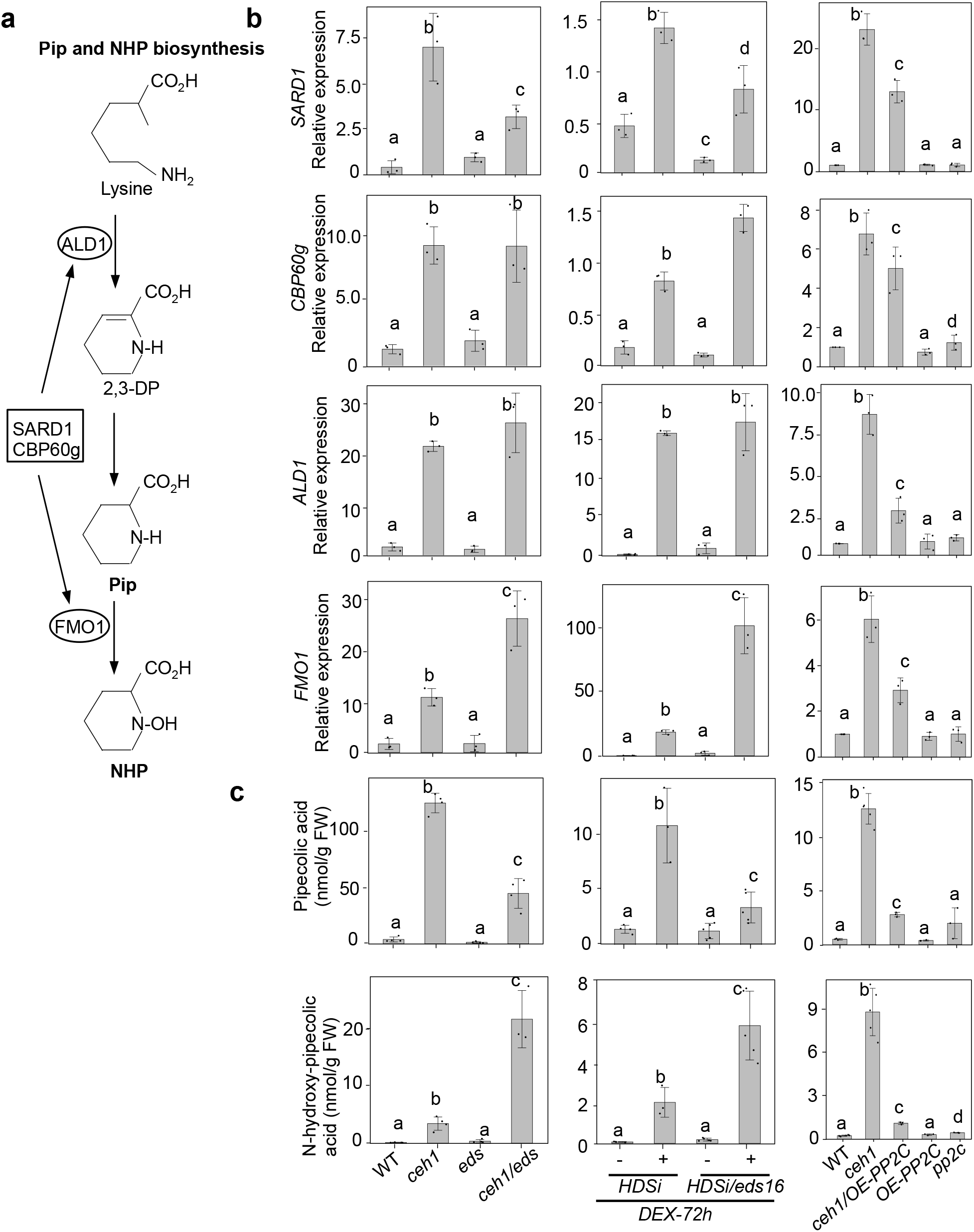
PP2C suppresses transcription of Pip and NHP biosynthesis genes and corresponding metabolites. (**a**) Schematic presentation of pipecolic acid (Pip) and N-hydroxy-pipecolic acid (NHP) biosynthesis pathway. (**b**) Relative expression levels of *SARD1*, *CBP60g*, *ALD1* and *FMO1* in WT, *ceh1, eds16, ceh1/eds16*, and in mock (-) and post 72 h DEX-treated *HDSi* and *HDSi/eds16* lines, and in *PP2C* overexpression in the wild-type (*OE-PP2C*) and the *ceh1* mutant (*ceh1/OE-PP2C*) backgrounds as well as in *pp2c* mutant, show reversion of SA- and MEcPP-dependent induction of these genes in *PP2C* overexpressing lines. (**c**) Measurements of Pip and NHP metabolites in aforementioned genotypes confirm their MEcPP- and SA-dependent alterations, and their reduced abundance in *PP2C* overexpression lines. All Data are mean ± SD of three biological and three technical replicates. Lower case letters on top of histograms represent statistically significant differences (*P* ≤ 0.05) by using ANOVA test.

Additional studies show that the overexpression of *PP2C* (*ceh1/OE-PP2C*) diminishes the MEcPP-mediated transcriptional induction of Pip and NHP biosynthetic genes, albeit at different degrees (Fig. **4b**). It is noteworthy that higher transcript levels of these genes in *ceh1* and *ceh1/OE-PP2C* relative to the WT, *OE-PP2C, or pp2C* may be due to the elevated MEcPP in the *ceh1* mutant. Lastly, similarly low expression levels of the genes in *pp2c* and WT lines could be attributed to the standard as opposed to stressed growth condition.

Next, we profiled Pip and NHP metabolite levels in aforementioned genotypes employed in the transcriptional profiling (Fig. **4c**). In concordance with the altered transcriptional profiles, accumulation of Pip and NHP metabolites is positively correlated to the presence of constitutively high or inducible MEcPP levels in *ceh1* and *HDSi* lines (Fig. **4c**). Whereas, the two metabolites are differentially accumulated in response to SA as evidenced by the reduced Pip content and enhanced NHP levels in *ceh1/eds16* relative to the *ceh1*mutant. In addition, decreased Pip and NHP levels in the *ceh1/OE-PP2C* relative to the *ceh1* mutant clearly support the PP2C-mediated reduction of both metabolites in spite of the high MEcPP levels.

Collectively, the finding establishes PP2C-mediated transcriptional suppression of Pip and NHP biosynthesis genes and by extension reduction of their corresponding metabolites, critical for eliciting systemic responses.

### PP2C interacts with MAPK3 and 6

To examine the subcellular site of PP2C action in high MEcPP containing *ceh1* mutant, we imaged the PP2C-GFP tagged *OE-PP2C* and *ceh1/OE-PP2C* lines, and confirmed plasma membrane, cytosolic and nuclear localization of the protein as previously reported for the WT background (Spartz *et al*., 2014; Tovar-Mendez *et al*., 2014; Ren *et al*., 2018) (Fig. **5a**).

**Figure 5.**
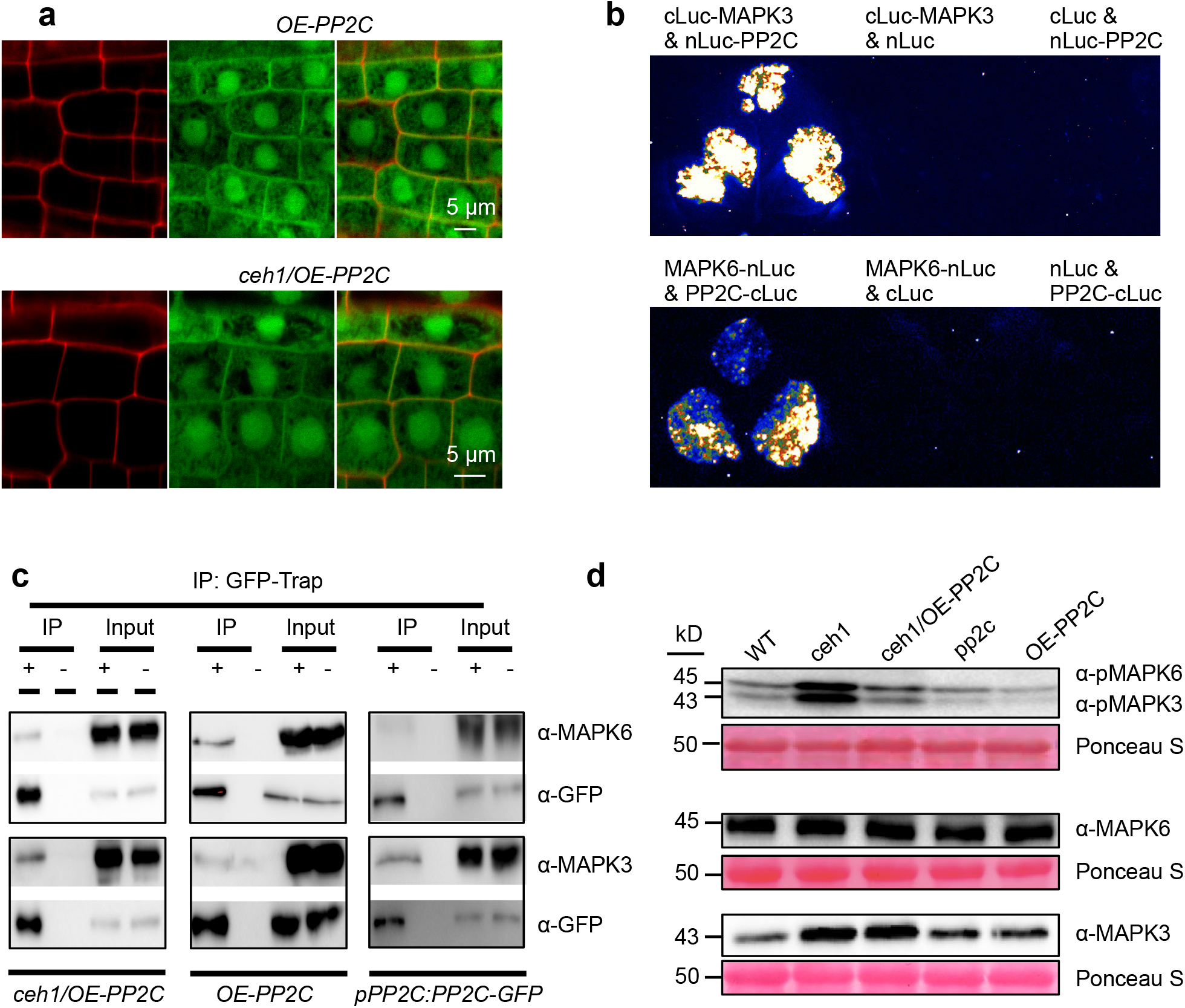
MAPK3/6 phosphorylation levels and their physical interaction with PP2C. (**a**) Confocal images of plasma membrane, cytosolic and nuclear localization of PP2C in the wild-type (*OE-PP2C*) and *ceh1* (*ceh1/OE-PP2C*) backgrounds overexpressing 35S::PP2C-GFP construct. (**b**) Representatives of split luciferase complementation assays in *Nicotiana benthamiana* displayed by dark-field images of leaves expressing cLuc-MAPK3 (C-terminal Luc fused with MAPK3) and nLuc-PP2C (N-terminal Luc fragment fused with PP2C) (upper panel) and MAPK6-nLuc (MAPK6 fused with N-terminal fragment of Luc) and PP2C-cLuc (PP2C fused with C-terminal fragment Luc) (lower panel). (**c**) The *in vivo* interaction of PP2C with MAPK3 and MAPK6 determined by co-immunoprecipitation assay. Protein samples obtained from *ceh1*/*OE-PP2C*, *OE-PP2C*, and *pPP2C:PP2C-GFP* seedlings grown under standard conditions were immunoprecipitated using GFP (+) and empty (-) magnetic beads. Immunoblots were analyzed using with α-MAPK3 or α-MAPK6. Each blot shows protein inputs before (input, right panels) and after (IP, left panels) immunoprecipitation. (**d**) Immunoblots show that *PP2C* overexpression in the *ceh1* mutant reverses the MEcPP-mediated high phosphorylation of MAPK3 and MAPK6 (α-pMAPK6 and α-pMAPK3, top panels), without notable impact on the protein abundance of MAPK6 (middle panels) or MAPK3 (bottom panels). Ponceau S staining shows protein loading.

To identify PP2C protein targets, we initially employed two independent methods. One method was based on immunoprecipitation-mass spectrometry (IP-MS) using a GFP specific antibody for IP of the PP2C interacting proteins in *ceh1/OE-PP2C* and *OE-PP2C* lines, and the control wild-type plant. As a second method, we employed a yeast-two-hybrid library-screening assay. The subsequent MS profiling of the samples derived from each of these two methods (Supplemental dataset 4), led to identification of several PP2C interacting proteins, most notably among them MAPK3 and 6.

Because of the indispensable function of these MAPKs in triggering Pip accumulation and by extension SAR induction (Wang *et al*., 2018), we verified their interactions with PP2C by additional methods. One method was based on the agro-infiltration-based transient assays in *Nicotiana benthamiana*. For this approach we used fusion constructs of MAPK3/6 and PP2C in various configurations (MAPK3 fused to carboxyl-terminal fragment of LUC, and PP2C fused to amino-terminal fragment of LUC, MAPK6 fused to amino-terminal fragment and PP2C fused to carboxyl-terminal fragment of LUC) (Figs. **5b** and **S7a-b**). The luciferase reconstitution-based activity is clearly evident in the leaves co-infiltrated with *PP2C* and *MAPK3* and *6* fusion constructs, but is absent in the leaves co-infiltrated with the respective controls. In a second independent approach, we examined the *in vivo* interaction of PP2C with MAPK3 and MAPK6 by targeted co-immunoprecipitation (CO-IP) using a GFP specific antibody for IP of PP2C-GFP in *ceh1/OE-PP2C, OE-PP2C* and *pPP2C:PP2C-GFP* lines, followed by immunoblot analyses using GFP as well as the MAPK3 and MAPK6 specific antibodies (Fig. **5c**). The clear and specific presence of an MAPK3 and MAPK6 reacting bands in the IP fractions of *ceh1*/*OE-PP2C* and *OE-PP2C* lines, but not in the control agarose beads, verified the *in vivo* interaction of PP2C with MAPK3 and MAPK6 proteins (Fig. **5c**).

Next, we explored the ramification of the interaction between MAPKs and PP2C protein by comparing the levels of phosphorylated MAPK3 and 6 in various genotypes (WT, *ceh1*, *ceh1/OE-PP2C, pp2c*, and *OE-PP2C*) using α-pMAPK6 and α-pMAPK3 antibodies deemed to specifically detect the respective phosphorylated proteins. The immunoblots clearly show similarly abundant phosphorylated kinases in WT and *pp2c*, however the levels are slightly but detectably reduced in *OE-PP2C* line grown under standard conditions (Fig. **5d**). Furthermore, these differences are not attributed to changes in the total MAPK3/6 protein abundance in WT, *pp2c* and *OE-PP2C* line, as they display similar levels on the immunoblots probed with the respective antibodies (Fig. **5d**). The most notable reduction in the abundance of phosphorylated MAPK3 and 6 is in *ceh1/OE-PP2C* compared to *ceh1* (Fig. **5d**). It is of note that the MAPK3 protein levels are similar between *ceh1* and *ceh1/OE-PP2C*, albeit more abundant than that of the other genotypes. This difference in abundance could contribute to higher levels of phosphorylated MAPK3 in the *ceh1* mutant relative to other genotypes, namely WT, *pp2c* and *OE-PP2C*. However, the higher level of phosphorylated MAPK3 in *ceh1* compared to *ceh1/OE-PP2C* is in spite of the similar abundance of total proteins in these genotypes. Additionally, altered levels of phosphorylated MAPK6 in various genotypes, most notably with heightened abundance in the *ceh1* mutant, is seemingly independent of slight variation of MAPK6 total protein levels amongst these genotypes (Fig. **5d**).

The above results collectively identify MAPK3 and 6 kinases as PP2C interacting proteins. Moreover, the inverse correlation between PP2C protein abundance and the phosphorylated levels of MAPK3/6 support the notion of PP2C function as the phosphatase.

### Abiotic and biotic stresses enhance MEcPP and induce Pip and NHP levels

We have previously established that the two most prevalent environmental insults, wounding and high-light induce MEcPP accumulation in plants (Xiao *et al*., 2012). This data together with MEcPP-mediated increased levels of Pip and NHP metabolites prompted us to examine the impact of mechanical damage and high-light on accumulation of these two SAR triggering metabolites. Thus, we examined the sequential steps of the events starting from MEcPP accumulation to the production of Pip and NHP in high-light treated and wounded seedlings.

We initially examined high-light treated plants and established their increased MEcPP and decreased auxin content using transgenic R2D2 reporter lines expressing auxin-degradable (DII) fluorescent protein as a proxy for IAA levels (Liao *et al*., 2015) (Figs **6a-b**). Reduced IAA content prompted us to examine the expression levels of *PP2C* and the auxin-responsive transcription factors, *ARF 7* and *19* in control and stressed seedlings (Fig. **6c**). Reduced expression levels of these genes in response to high-light treatment are reminiscent of our earlier observation in the *ceh1* mutant (Fig. **2c**). Reduced *ARF 7* and *19* transcript levels in stressed seedlings prompted us to examine the ramification of these reductions on the expression of genes within Pip and NHP biosynthetic pathway. As such, we compared the transcript levels of *CBP60, ALD1* and *FMO1* in the WT and *arf7/19* mutant lines (Fig. **S8**). The clear enhancement of the transcript levels of all genes in the *arf7/19* mutant line alludes to the suppressive function of ARF9 and 17 on the expression of Pip and NHP biosynthetic-pathway genes. The result provides a rationale for the stress-mediated suppression of ARF7 and 19, hence enabling increased production of Pip and NHP.

**Figure 6.**
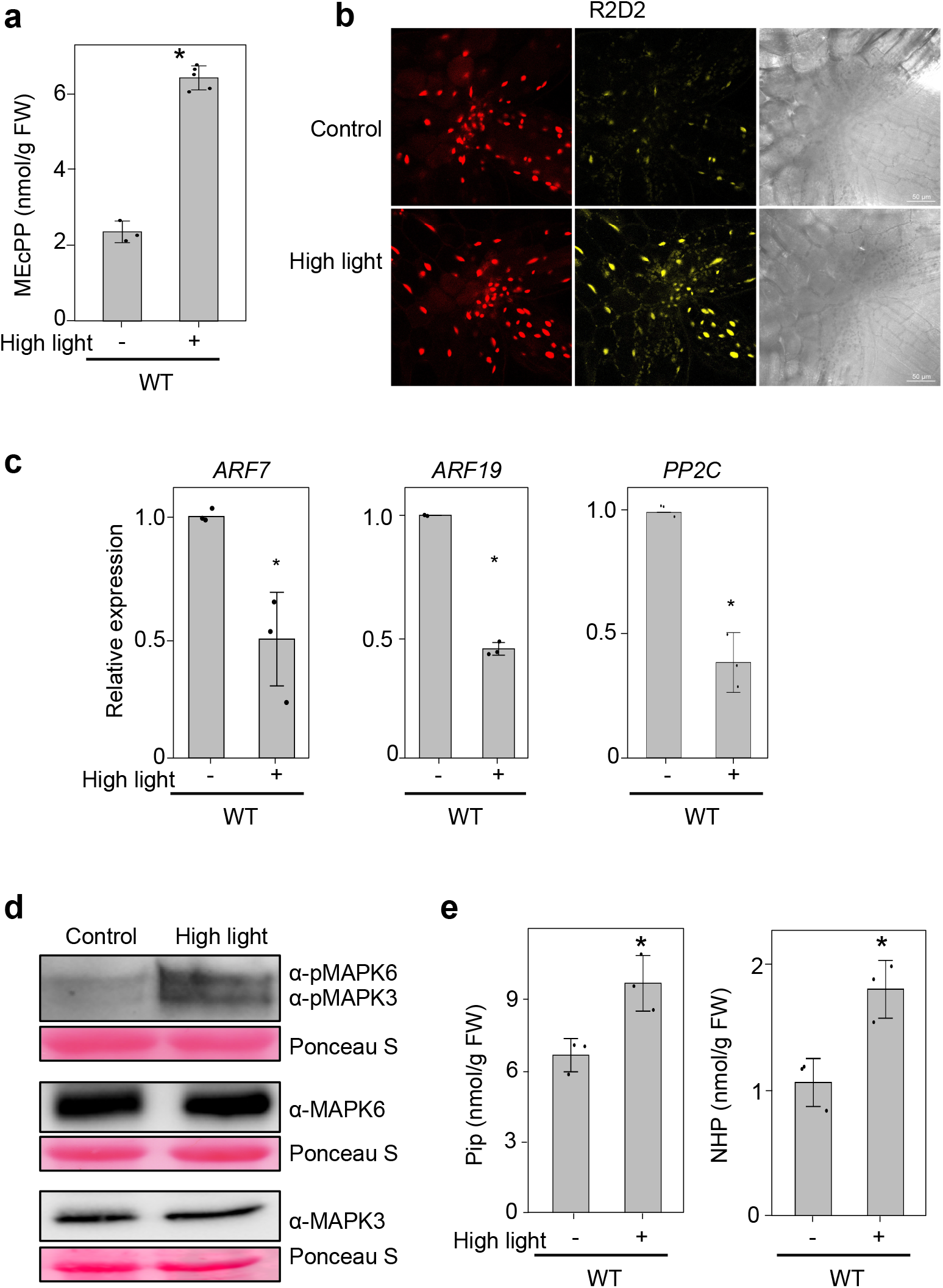
High light induces MEcPP accumulation and enhances Pip and NHP abundance. (**a**) Induction of MEcPP levels and (**b**) reduction of IAA abundance 90 min post high light (800 μmol m^-2^s^-1^, 90 min) treatment as measured by fluorescence of mDII-ntdTomato/DII-n3xVenus and the corresponding bright-field images of control and high light-treated auxin reporter R2D2 seedlings. (**c**) Relative expression levels of *ARF 7/19* and *PP2C* in high-light-treated seedlings. (**d**) Immunoblot analyses of total and phosphorylated MAPK3/6 in high-light-treated seedlings. Ponceau S show equal protein loading. (**e**) Enhanced levels of Pip and NHP metabolites in high light-treated seedlings. Two-tailed Student’s *t* tests and ANOVA tests are used for the statistical analyses and the asterisk and different letters denote significance (*P* ≤ 0.05).

Next, we examined the relative abundance of phosphorylated MAPK3 and 6 in the high-light treated seedlings compared to the control (Fig. **6d**). The data establishes similar abundance of their total proteins but enhanced levels of phosphorylated MAPK3/6 under high light condition, (Fig. **6d**). Next we compared the abundance of Pip and NHP metabolites in the stressed versus the control seedlings and confirmed high-light-induced accumulation of the metabolites (Fig. **6e**). The finding supports the notion of Pip and NHP triggering SAA in response to high-light exposure.

Subsequently, we extended these studies to include unwounded and wounded (90 min post wounding) *pp2c* mutant and wild-type plants. The finding clearly establishes wound-induced MEcPP-accumulation independently of PP2C (Fig. **7a**). Furthermore, similarly to high-light-treated plants, auxin levels as well as the expression levels of *ARF7, 19* and *PP2C* are reduced in wounded relative to unwounded plants (Fig. **7b-c**). Moreover, wounding induces phosphorylation of MAPK3 and 6 in the WT and *pp2c* mutant (Figs. **7d** **and S9**). Accordingly, there is increased accumulation of Pip and NHP in both backgrounds albeit at higher levels in *pp2c* compared to the WT (Fig. **7e**).

**Figure 7.**
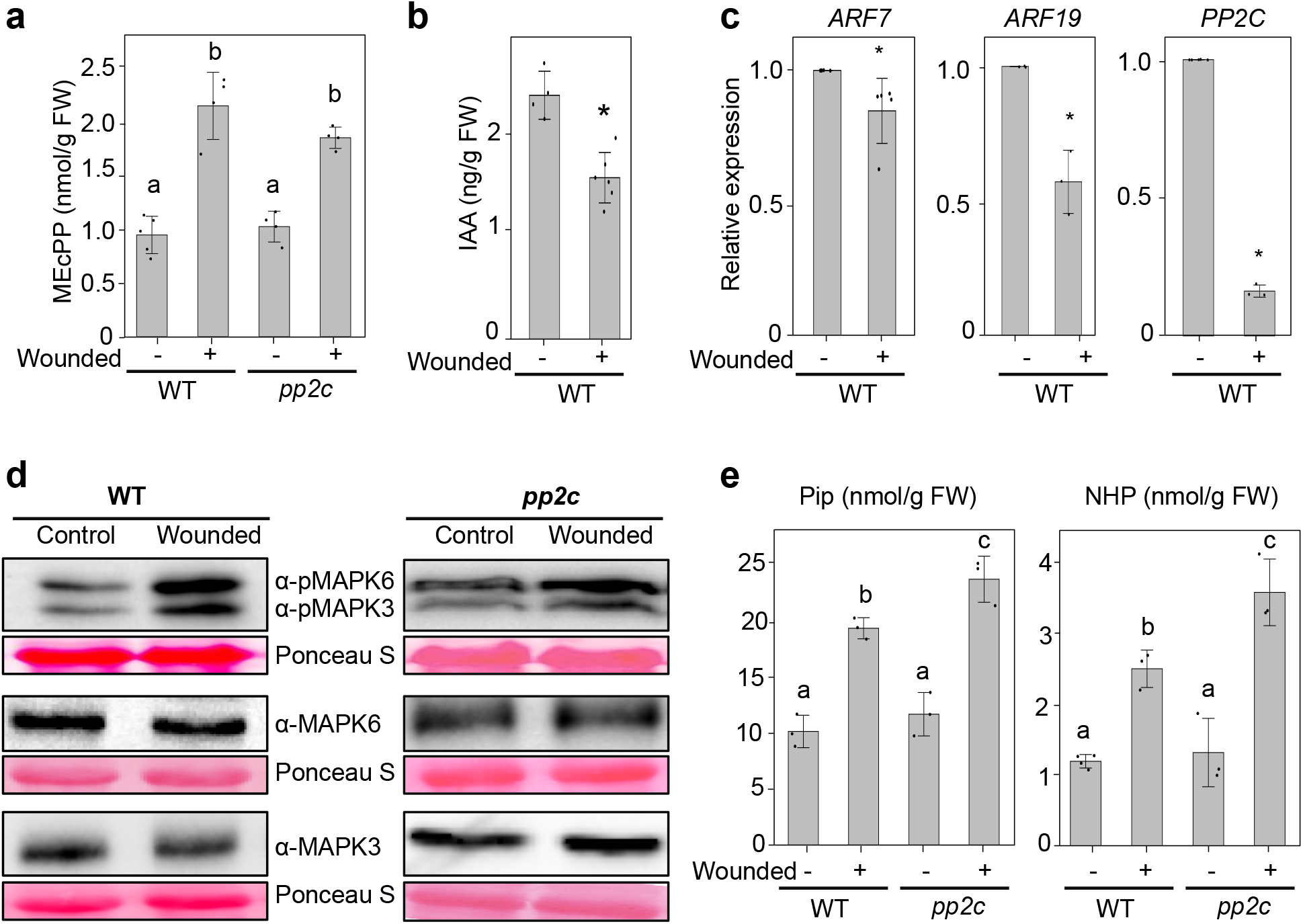
Wounding induces MEcPP accumulation and enhances Pip and NHP abundance. (**a**) Induction of MEcPP levels and (**b**) reduction of IAA abundance in 90 min post wounded WT and *pp2c* mutant seedlings compared to control unwounded plants. (**c**) Relative expression levels of *ARF 7/19* and *PP2C* in unwounded and wounded WT plants. (**d**) Immunoblot analyses of total and phosphorylated MAPK3/6 in unwounded and wounded WT and *pp2c* mutant seedlings. Ponceau S show equal protein loading. (**e**) Metabolic analyses of Pip and NHP in wounded and WT and *pp2c* mutant seedlings. Two-tailed Student’s *t*-tests and ANOVA tests are used for the statistical analyses and the asterisk and different letters denote statistical significance (*P* ≤ 0.05).

Next we examined the multi-component signaling cascade potentiating the biosynthesis of SSR in plants challenged with two biotic stresses, aphids (*Macrosiphum euphorbiae*) and a viral pathogen [*Cucumber mosaic virus* (*CMV-m2b*)] (Fig. **8a-d**). The data clearly show increased MEcPP levels upon aphid infestation and CMV infection, followed by decreased expression of *ARF7/19* and *PP2C*, and the ensued heightened MAPK3 and 6 phosphorylation albeit without altered abundance of the total proteins, and finally induction of Pip and NHP metabolites.

**Figure 8.**
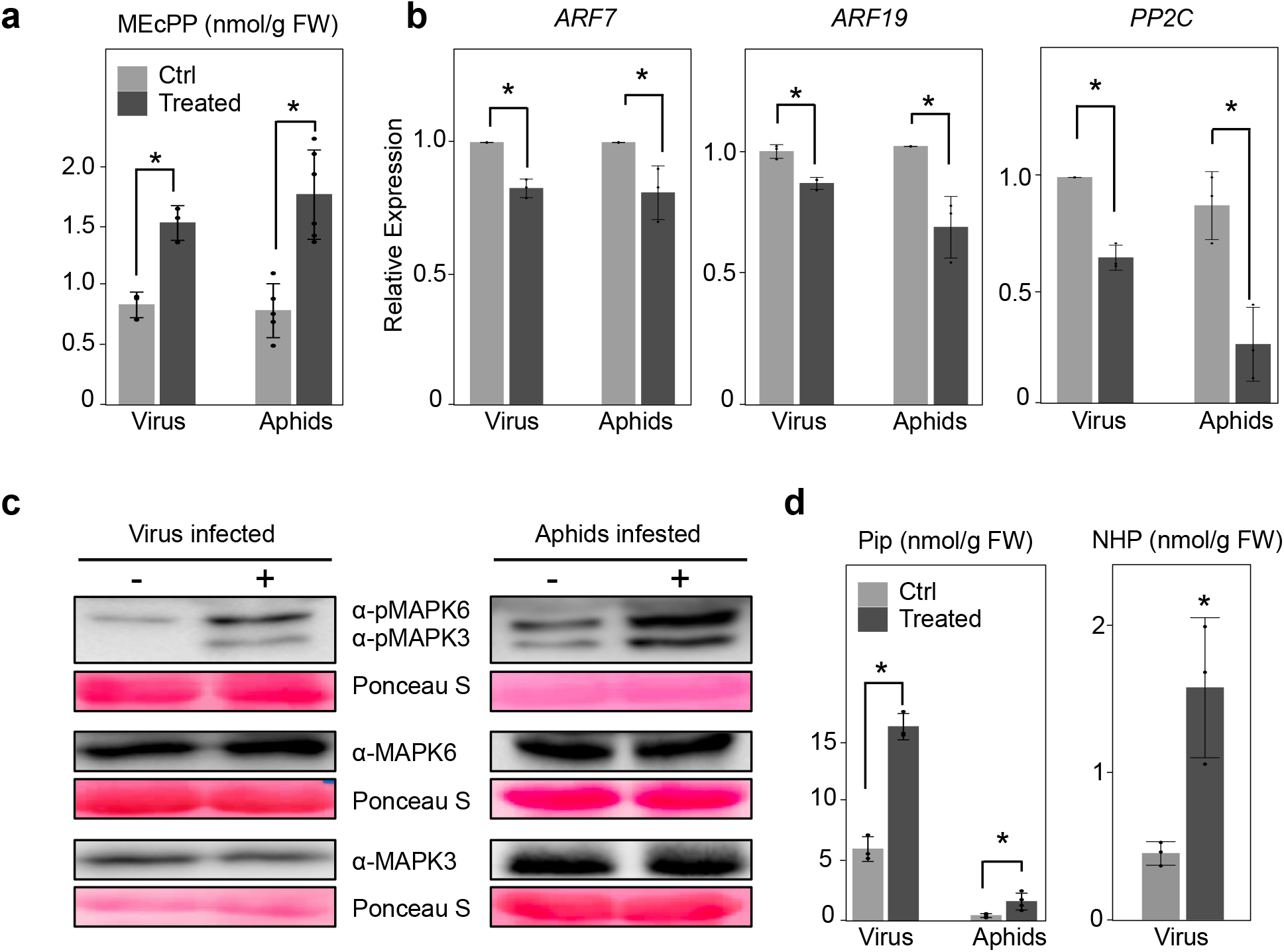
Aphid infestation and viral infection induce MEcPP accumulation and enhance Pip and NHP abundance. (**a**) Induction of MEcPP post 2 weeks of cucumber mosaic virus (CMV-M2b) infection, and 24h aphid infestation in WT seedlings, and (**b**) suppression of *ARF 7/19* and *PP2C* expression in biotically challenged plants. (**c**) Immunoblot analyses of total and phosphorylated MAPK3/6 in mock (-) and virus/aphids (+) treated seedlings. Ponceau S show equal protein loading. (**d**) Metabolic analyses of Pip and NHP in mock (-) and virus/aphids (+) treated seedlings. The asterisk denotes statistical significance (*P* ≤ 0.05) by using two-tailed Student’s *t*-tests.

Collectively, our findings establish a link between the two prevalent naturally occurring abiotic stresses and biotic insults to the induction of MEcPP levels followed by the reduction in auxin abundance and the consequential decline in *PP2C* expression, and ultimately phosphorylation of MAPK3/6 and accumulation of Pip and NHP metabolites. As such, the data supports functional expansion of Pip and NHP in eliciting SSR by abiotic or biotic triggers.

## Discussion

Plants have evolved complex tiers of molecular and biochemical networks to detect, transmit and amplify adaptive signals for dynamic restoration of cellular homeostasis and function at the local and the distal site of insults. This broad spectrum response known as systemic stress response (SSR) appears to be conserved in plants across the plant kingdom, and as such the focus of intense studies (Shah & Zeier, 2013). However, the identity of signals that initiate this response has remained fragmentary. Here, we provide a complete module of the nature and the sequence of events that trigger SSR cascade (Fig. **9**). Specifically, we illustrate that accumulation of the stress-specific plastidial retrograde signaling metabolite, MEcPP, achieved either by genetic manipulation or via challenging plants with the two prevalent abiotic challenges (mechanical damage and high-light treatment) and two biotic insults (aphids and CMV), result in reduced auxin concentration and the consequential decreased expression of *ARF7* and *19*, the transcriptional activators of *PP2C*. Auxin-induction of *PP2C* corroborates the earlier finding (Nemhauser *et al*., 2006) and supports the notion of PP2C function as a suppressor of stress-response genes and an inducer of the growth-related genes. Although the notion is contradicted by the report of auxin-induced SMALL AUXIN UP-RNA binding to, and inhibiting SAUR19. This results in inactivation of PP2C and consequential phosphorylation and hence activity of H+-ATPase required for cell expansion in the apical hook of etiolated seedlings, where PP2C is abundantly present (Spartz *et al*., 2014; Ren *et al*., 2018). Indeed, induction of *PP2C* expression by auxin and suppression of its enzyme activity by the auxin-induced SAUR19 represent two conflicting auxin-based regulatory responses. These opposing responses are prime examples of multi-layered auxin-based fine-tuning of PP2C both at the expression and at the enzyme activity levels. This delicate balance shifts the function of PP2C enzyme to either a positive or a negative regulator of growth, depending upon the tissue, and tailored to nature of the environmental challenges.

**Figure 9.**
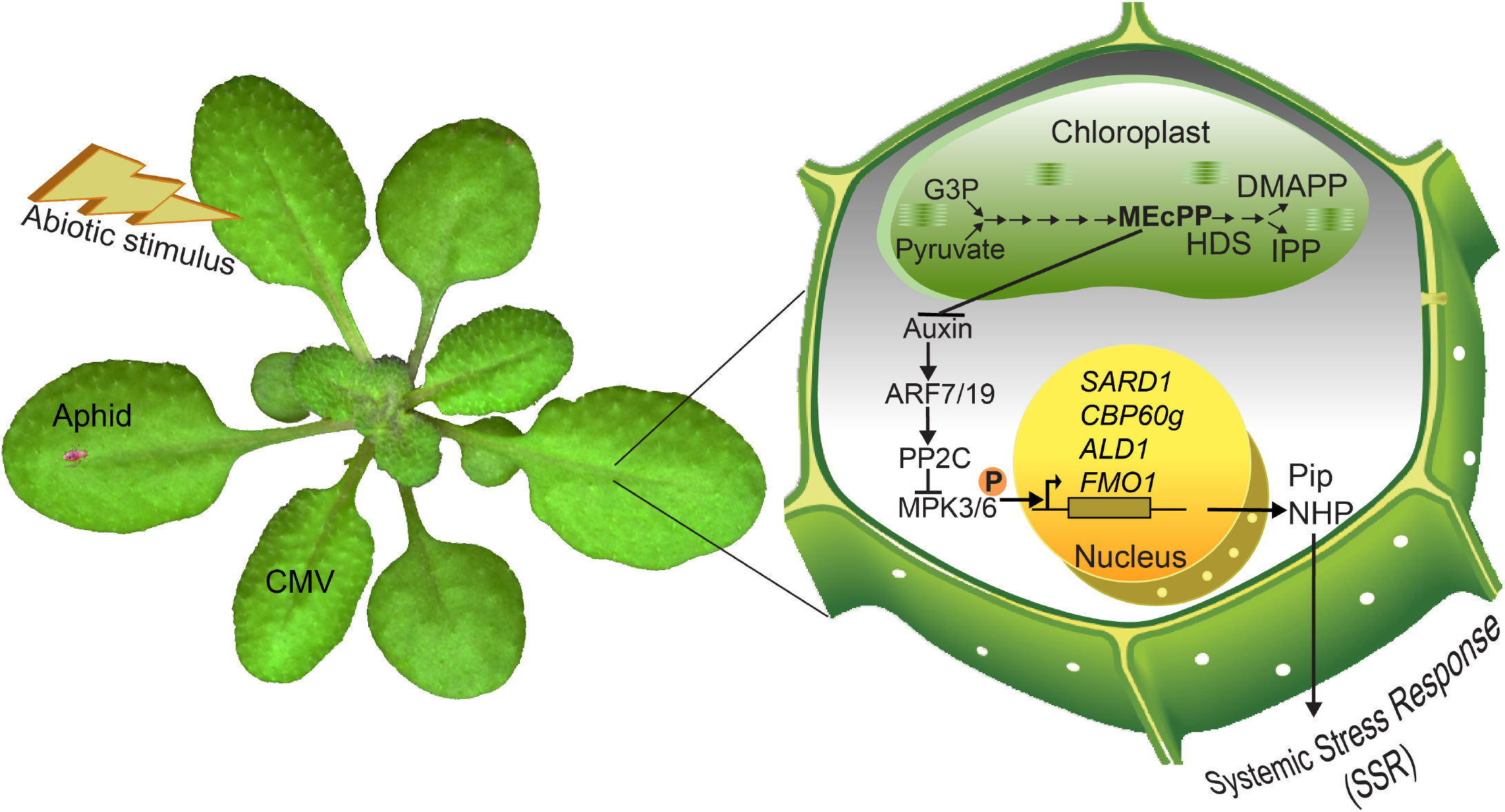
Biotic and abiotic insults trigger the retrograde signaling cascade initiating SSR. Schematic model depicting biotically and abiotically-induced MEcPP-accumulation mediates reduction of auxin abundance that lessens expression levels of the ARF7/19, the transcriptional activators of *PP2C*. This enables phosphorylation of MAPK3/6 required for induction of Pip and NPH biosynthesis genes and production of their respective metabolites key to activation of general SSR.

Combined parallel and independent approaches of IP-MS, yeast-two-hybrid library screening, split-luciferase assay and CO-IP followed by immuno-blot analyses verified physical interactions between MAPK3/6 and PP2C protein. The biological ramification of this interaction is best captured by enhanced levels of phosphorylated MAPK3 and 6 in the constitutively (*ceh1* mutant) and stress-inducible MEcPP accumulating plants, and conversely by their reduced phosphorylated forms in *ceh1/OE-PP2C* lines, raising the likelihood of PP2C function as the responsible phosphatase. This notion is supported by the reduction of MAPK3/6 phosphorylation in *OE-PP2C* line compared to WT. It is of note that the lack of enhanced MAPK3/6 phosphorylation in *pp2c* mutant line under standard condition, is likely due to the absence of a stress-activated kinase within the MAPK kinase kinases cascade, otherwise present in MEcPP accumulating *ceh1* mutant, and in biotically and abiotically challenged plants. This scenario is supported by H_2_O_2_ activation of ANP1, an *Arabidopsis* MAPKKK that initiates a phosphorylation cascade including MAPK3/6 followed by induction of specific stress-responsive genes, and suppression of auxin-inducible genes (Kovtun *et al*., 2000). Indeed, the functional input of MAPKs in mis-localizing polar auxin transport proteins (PINs) expands the regulatory roles of activated MAPKs in hampering auxin distribution and signal transduction (Jia *et al*.,2016; Dory *et al*., 2018). This process may therefore constitute the underlying mechanism of reduced abundance of PIN1 in the MEcPP accumulating *ceh1* mutant, where MAPK phosphorylation of PIN1 result in mis-localization and degradation of this auxin transporter (Jiang *et al*., 2018). If so, this places the reduction of *PP2C* transcript and the consequential activation of MAPK3/6 at the interface between MEcPP and auxin signaling, and uncovers the sequence of events between the two regulatory capacities required for tailoring plant growth and developmental responses to environmental cues.

The converse correlation between PP2C abundance and the prevalence of SAR inducible genes in MEcPP-accumulating plants, confirmed by qRT-PCR analyses of targeted genes involved in Pip and NHP biosynthesis genes, is in agreement with the increase of the respective metabolite levels. Furthermore, analyses of constitutive and inducible MEcPP accumulating plant that are either deficient in, or contain high SA, show differential SA-mediated transcriptional responses. That is, except for the *SARD1* whose induction is partly dependent on SA, the expression levels of the other three genes are either SA-independent (*ALD1*) or suppressed by SA (*CBP60g* and *FMO1*). In contrast, MEcPP induces the expression of all these genes albeit at different degrees, placing *SARD1* as the least and *FMO1* as the most MEcPP-responsive genes. Moreover, analyses of Pip and NHP metabolite levels show that while SA induces Pip production, it suppresses NHP levels, whereas MEcPP-mediates induction of both metabolites although at different levels. This establishes MEcPP as the inducer of SSR, and further suggests that despite the critical role of SA as a mobile signal for SAR (Neuenschwander *et al*., 1995; Park *et al*., 2007), SA is not essential for production of Pip and NHP.

Additionally, increased levels of Pip and NHP in plants challenged with wounding and high-light expands the role of these SAR triggering metabolites to the establishment of a resistance state not only when confronted with biotic but also when challenged with abiotic insults as evidenced by the reported *FMO1* induction in response to H_2_O_2_ accumulation (Chen & Umeda, 2015).

In summary, here we reveal the nature and the organization of a multicomponent retrograde signaling cascade that induces the biosynthesis of Pip and NHP in response to both biotic and abiotic insults. This occurs through alterations of positive and negative regulators that enable a timely modification of expression profiles and the consequential reconfiguration of the metabolic network for optimal implementation of this adaptive response as a general strategy to fend against a complex myriad of insults. We specifically identify plastids as the initiation site and the plastidial retrograde signaling metabolite, MEcPP, as the initiating signal potentiating the concerted arrays of signaling network responsible for production of Pip and NHP, the triggers of systemic stress responses in face of a myriad of environmental challenges.

## Supporting information

Supplemental data sets

## Acknowledgements

We would like to thank Professor Elizabeth Sattely at Stanford University for providing us with the chemically synthesized NHP standard. We would like to thank our colleagues at UCR, Prof. Isgouhi Kaloshian and Dr. Jacob Macwilliams for aphid-infested plants, and Dr. Bailong Zhang for viral-infected plants. This work was supported by NSF CREATE-IGERT training program (NSF DGE-0653984) and NSF-GRFP 1148897 to M.L, by Dr. John W. Leibacher and Mrs. Kathy Cookson endowed chair funds to KD and by National Institutes of Health (NIH) GM067203 to W.M.G., and by NIH, R01GM107311-8 grant to KD.

## Authors’ contributions

L. Z., J.Z.W. and KD designed the study, L.Z., J.Z.W., X.H., H. K. and M. L. performed the experiments, W. M. G. provided antibody and K.D. wrote the manuscript.

## Supporting Information

**Figure S1. Reduced expression of *PP2C* in the *ceh1* mutant**

(**a**) Phylogeny of PP2C family members in clade D. (**b**) RNA-seq-based analyses of relative expression levels of *PP2C.D* family members show decreased levels of *PP2C.D1* and *PP2C.D7*, and increased levels of *PP2C.D8* and *PP2C.D9* expression in the *ceh1* mutant relative to the wild-type plant.

**Figure S2. PP2C immunoblot.**

(**a**) Detection of PP2C protein using PP2C antibody on an immunoblot of protein extracts from various genotypes. (**b**) Ponseau S stain shows the equal loading.

**Figure S3 Relative expression levels of selected *ARF* family members**

Total RNAs isolated from two-week-old wild-type (WT), *ceh1/eds16*, *eds16* seedlings were subjected to q-PCR analyses. Relative expression levels of representative *ARFs* (ARF2, *ARF3, ARF10, ARF11* and *ARF18*) were normalized to the levels of *At4g26410* (M3E9). All Data are mean ± SD of three biological and three technical replicates. Two-tailed Student’s *t* test confirms MEcPP-independent expression of these *ARF* members.

**Figure S4. GO term analyses implicate PP2C as a growth-promoter and a stress-suppressor**

Comparative GO term analyses of induced genes in the *pp2c* mutant and *PP2C* overexpressing line (*OE-PP2C*) implicate PP2C as a stress suppressor and a growth promoter. The red bar shows the -Log_10_ P-values of altered transcript levels.

**Figure S5. PP2C is likely involved in biotic stress responses.**

(**a**) KEGG pathway enrichment analyses of induced genes in the *pp2c* mutant compared to Col, implicating PP2C as a biotic stress suppressor. The red bar shows the -Log_10_ P-values of enriched pathways. (**b**) The expression level of *PP2C* is significantly suppressed by pathogen-associated molecular patterns (PAMPs) treatment (flg22 and nlp20) of Col post 90 and 180 min. Data retrieved from the recently published report (Bjornson *et al*., 2021). The star on each histogram indicates significant changes of the expression level compared to the representative mock treatment.

**Figure S6. PP2C overexpression modifies transcriptional profile**

Venn diagrams illustrate significantly reduced number of SAR-induced (**a**) and Pip-induced genes (**b**) in *PP2C* overexpressing *ceh1* and inducible *HDSi* lines.

**Figure S7. Split luciferase complementation assays in *Nicotiana benthamiana***

Representatives of split luciferase complementation assays in *Nicotiana benthamiana* displayed by bright-field images of leaves expressing cLuc-MAPK3 (C-terminal Luc fused with MAPK3) and nLuc-PP2C (N-terminal Luc fragment fused with PP2C) (upper panel) and MAPK6-nLuc (MAPK6 fused with N-terminal fragment of Luc) and PP2C-cLuc (PP2C fused with C-terminal fragment Luc) (lower panel). Negative controls for each constructs include cLuc-MAPK3 & nLuc; cLuc & nLuc-PP2C; MAPK6-nLuc & cLuc and nLuc & PP2C-cLuc.

**Figure S8. Induction of *CBP60g, ALD1* and *FMO1* transcript levels in *arf7/19* mutant.**

Total RNAs isolated from two-week-old wild-type and *arf7/19* double mutant seedlings were subjected to q-PCR analyses. Relative expression levels of *CBP60g*, *ALD1* and *FMO1* were normalized to the levels of *At4g26410* (M3E9). All Data are mean ± SD of three biological and three technical replicates. The star represents the significantly statistic differences by two-tailed Student’s *t* test.

**Figure S9. Normalized relative intensity of protein levels.**

Phosphorylated MAPK6 (pMAPK6) and MAPK3 (pMAPK3), and un-phosphorylated MAPK6 and MAPK3 proteins are normalized to the levels of Ponceau stain of Rubisco.

**Table S1. Percentage of MEcPP-induced and PP2C-supppressed SAR- and Pip-inducible genes, and the RSRE containing genes**

**Table S2. GO term analyses of the up-regulated genes in *pp2c* mutant and *OE-PP2C*.** The presented number is -Log10 (P-value) for each of the GO term.

**Table S3. List of used primers**

**Supplemental data sets 1-3. List of differentially expressed genes**

**Supplemental data set 4. List of identified proteins in IP-MS**

