## Supplemental Figures for "A plastidial retrograde-signal potentiates biosynthesis of systemic stress response activators"

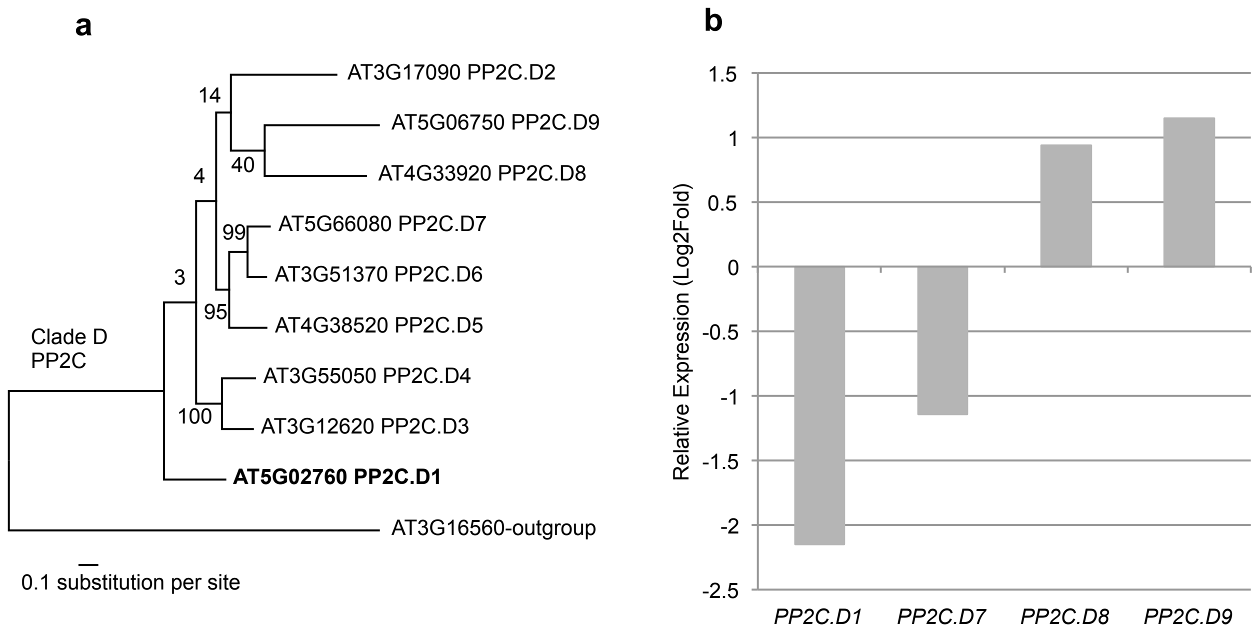


**Figure S1. Reduced expression of *PP2C* in the *ceh1* mutant.**
(**a**) Phylogeny of PP2C family members in clade D. (**b**) RNA-seq-based analyses of relative expression levels of *PP2C.D* family members show decreased levels of *PP2C.D1* and *PP2C.D7*, and increased levels of *PP2C.D8* and *PP2C.D9* expression in the *ceh1* mutant relative to the wild-type plant.

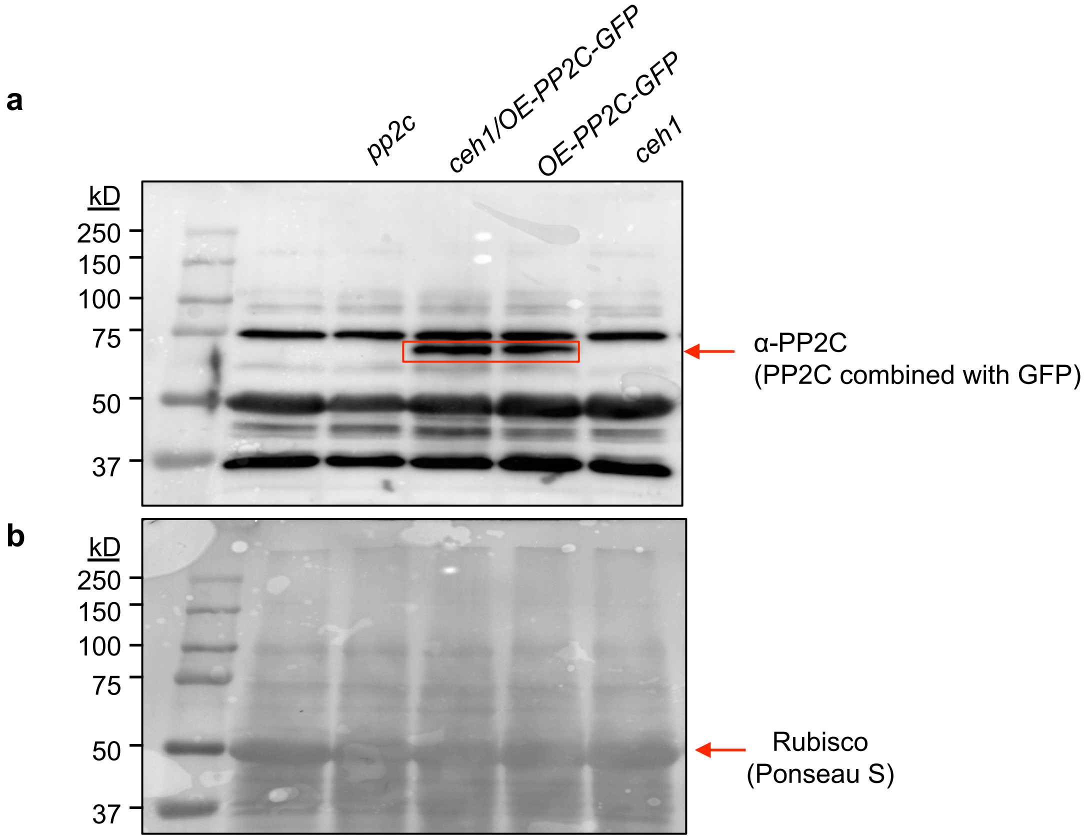


**Figure S2. PP2C immunoblot.**
(**a**) Identification of PP2C protein using PP2C antibody on an immunoblot of protein extracts from various genotypes.
(**b**) Ponseau S stain shows the equal loading.

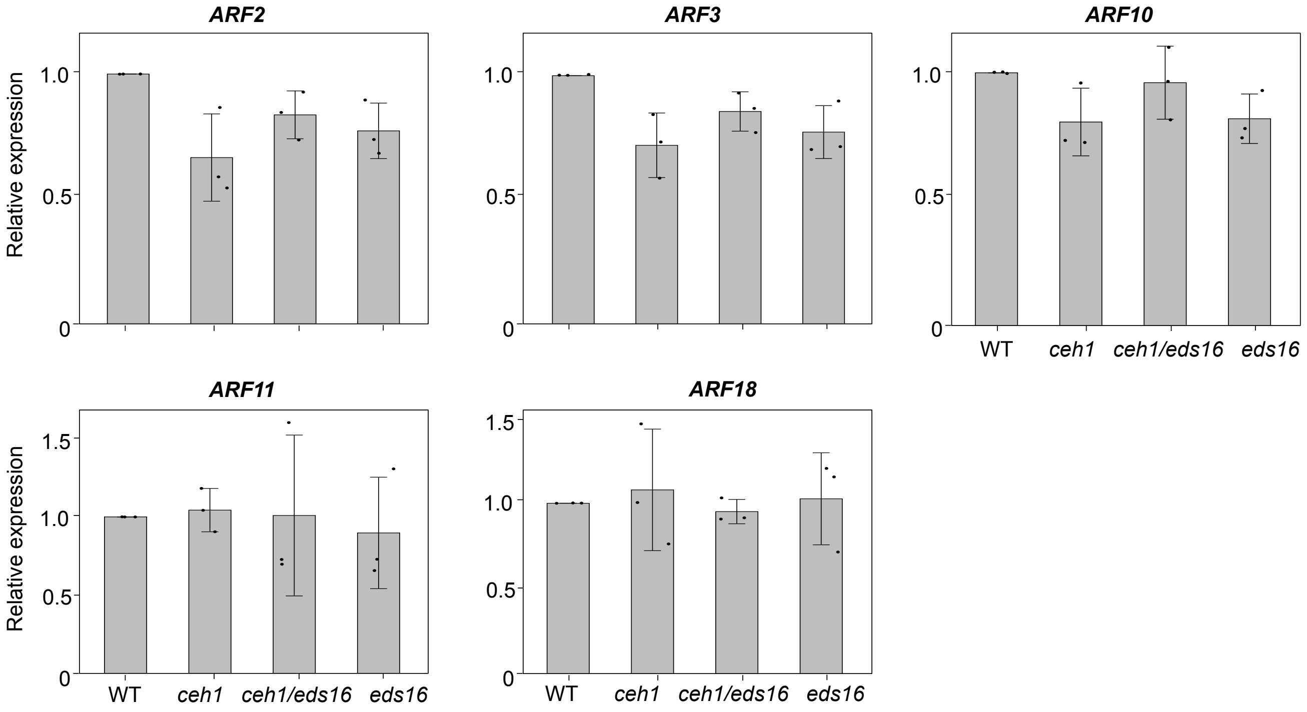


**Figure S3. Relative expression levels of selected *ARF* family members.**
Total RNAs isolated from two-week-old wild type (WT), *ceh1/eds16*, *eds16* seedlings were subjected to q-PCR analyses. Relative expression levels of representative *ARF*s (ARF2, *ARF3*, *ARF10*, *ARF11* and *ARF18*) were normalized to the levels of *At4g26410* (M3E9). All Data are mean ± SD of three biological and three technical replicates. Two-tailed Student's *t* tests confirms MEcPP-independent expression of these *ARF* members.

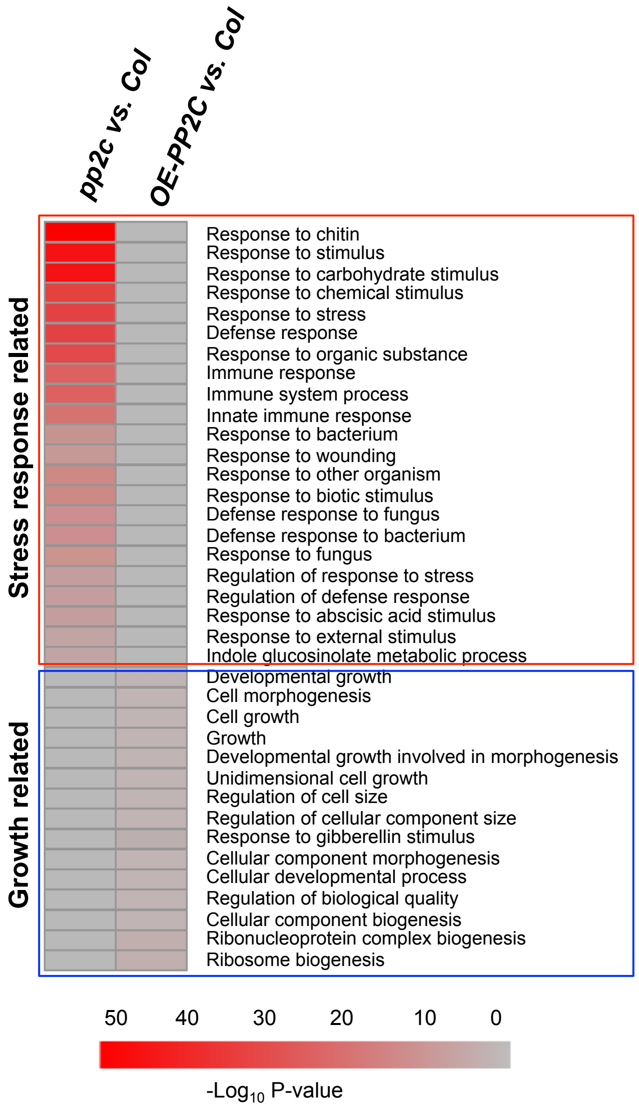


**Figure S4. GO term analyses implicate PP2C as a growth-promoter and a stress-suppressor.** Comparative GO term analyses of induced genes in the *pp2c* mutant and *PP2C* overexpressing line (*OE-PP2C)* implicate PP2C.D1 as a stress suppressor and a growth promoter. The red bar shows the $-Log_{10}$ P-values of enriched GO terms.

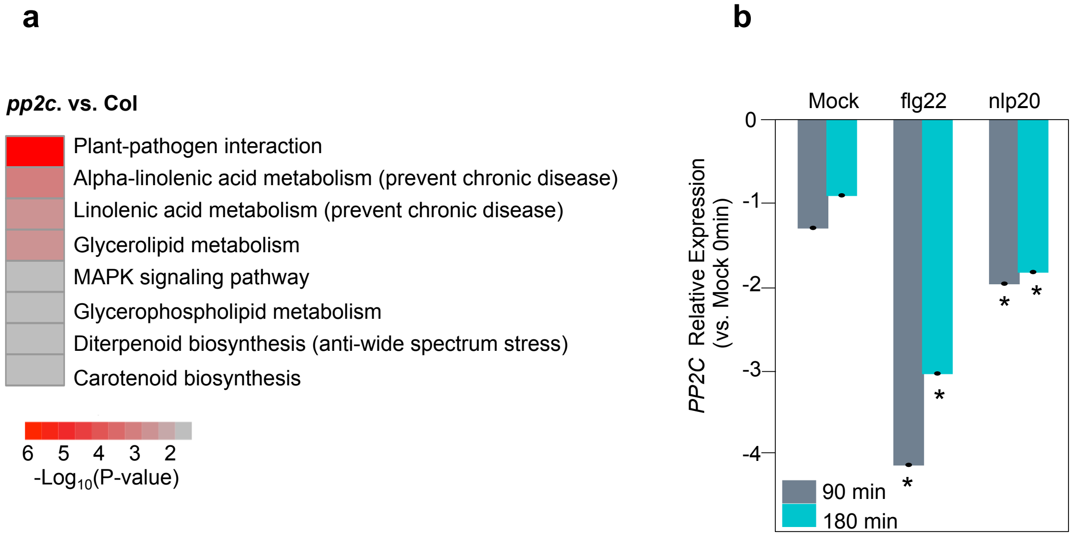


**Figure S5. PP2C is likely involved in biotic stress response.**
(**a**) KEGG pathway enrichment analyses of induced genes in the *pp2c* mutant compared to Col, implicating PP2C as a biotic stress suppressor. The red bar shows the -$Log_{10}$ P-values of enriched pathways.
(**b**) The expression level of *PP2C* is significantly suppressed by pathogen-associated molecular patterns (PAMPs) treatment (flg22 and nlp20) of Col post 90 and 180 min. Data retrieved from the recently published report (Bjornson et al., 2021). The star on each histogram indicates significant changes of the expression level compared to the representative mock treatment.

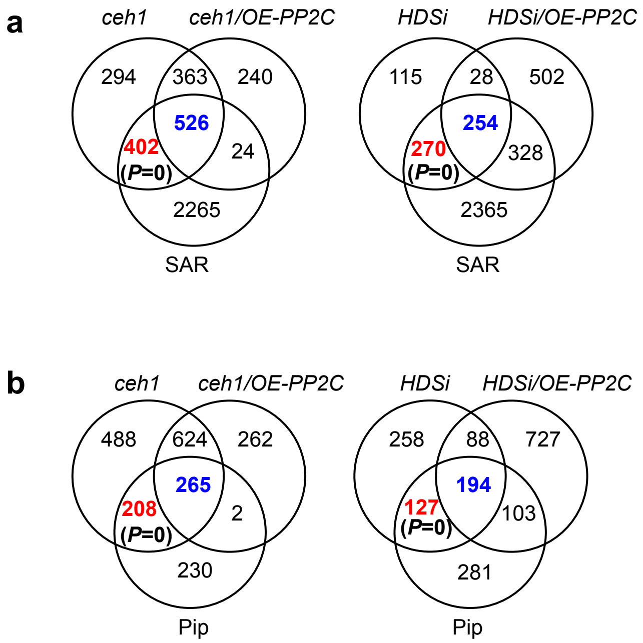


**Figure S6. PP2C overexpression modifies transcriptional profile**
Venn diagrams illustrate significantly reduced number of SAR-induced (**a**) and Pip-induced genes (**b**) in *PP2C* overexpressing *ceh1* and inducible *HDSi* lines.

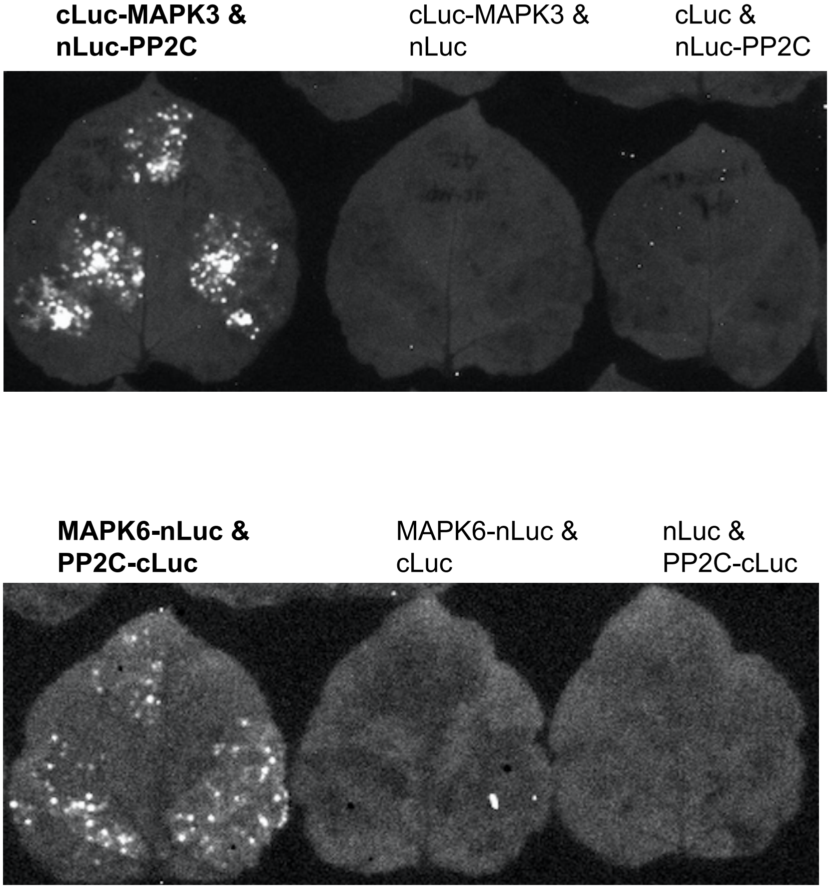


**Figure S7. Split luciferase complementation assays in *Nicotiana benthamiana.***
Representatives of split luciferase complementation assays in *Nicotiana benthamiana* displayed by bright-field images of leaves expressing cLuc-MAPK3 (C-terminal Luc fused with MAPK3) and nLuc-PP2C (N-terminal Luc fragment fused with PP2C) (upper panel) and MAPK6-nLuc (MAPK6 fused with N-terminal fragment of Luc) and PP2C-cLuc (PP2C fused with C-terminal fragment Luc) (lower panel). Negative controls for each constructs include cLuc-MAPK3 & nLuc; cLuc & nLuc-PP2C; MAPK6-nLuc & cLuc and nLuc & PP2C-cLuc.

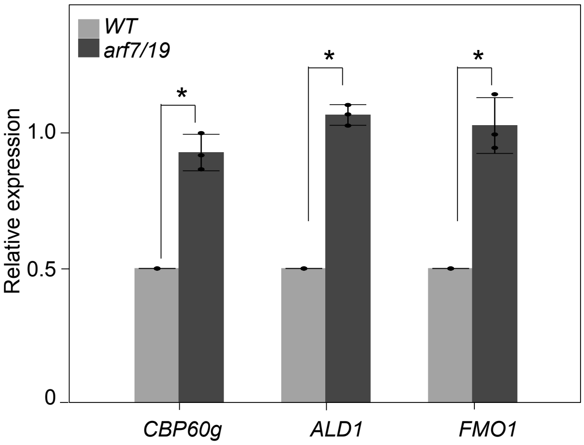


**Figure S8. Induction of *CBP60g*, *ALD1* and *FMO1* transcript levels in *arf7/19* mutant.**
Total RNAs isolated from two-week-old wild type (WT) and *arf7/19* double mutant seedlings were subjected to q-PCR analyses. Relative expression levels of *CBP60g*, *ALD1* and *FMO1* were normalized to the levels of *At4g26410* (M3E9). All Data are mean ± SD of three biological and three technical replicates. The star represents the significantly statistic differences by two-tailed Student's *t* test.

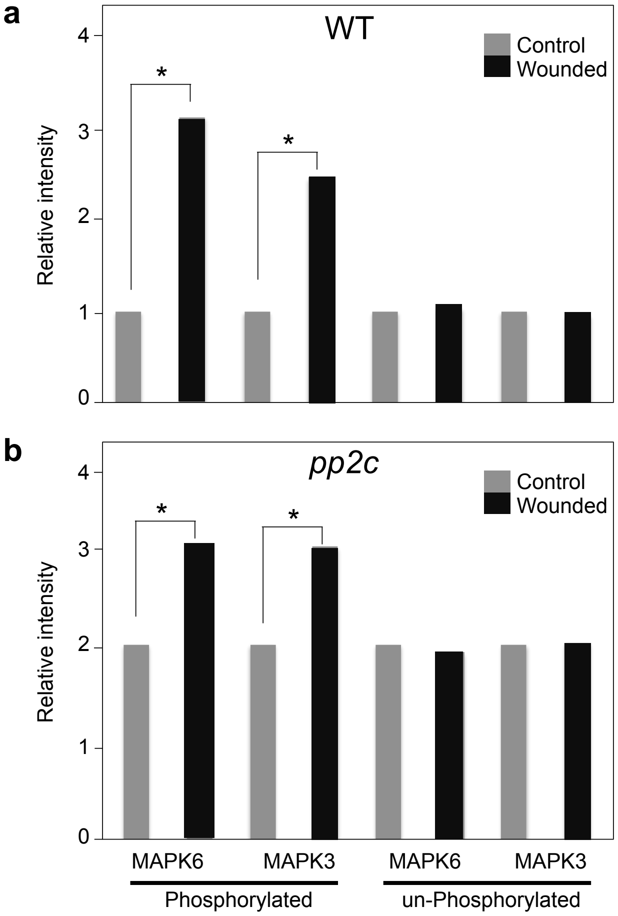


**Figure S9. Normalized relative intensity of protein levels.**
Normalized phosphorylated MAPK6 (pMAPK6) and MAPK3 (pMAPK3), and un-phosphorylated MAPK6 and MAPK3 proteins are normalized to the levels of Ponceau stain of Rubisco.
