## Supplemental Tables for "A plastidial retrograde-signal potentiates biosynthesis of systemic stress response activators"

**Table S1. Percentage of MEcPP-inducible and PP2C-suppressed SAR and Pip genes.**

|  | ***ceh1* background** | ***HDSi* background** |
| --- | --- | --- |
| Percentage of **RSRE**-motif containing genes in the *PP2C*-suppressed MEcPP-induced-genes | 47%  (Fig 3a, left panel) | 68%  (Fig 3a, right panel) |
| Percentage of MEcPP-induced-**SAR**-genes suppressed by *PP2C* | 43%  (Fig S4a, left panel) | 52%  (Fig S4a, right panel) |
| Percentage of MEcPP-induced-**Pip**-genes suppressed by *PP2C* | 44%  (Fig S4b, left panel) | 40%  (Fig S4b, right panel) |

**Table S2. GO term analyses by using the up-regulated genes in *pp2c* mutant and *OE-PP2C*.** The presented number is -Log10 (P-value) of each GO term.

|  | **GO terms** | ***pp2c* mutant** | ***OE-PP2C*** |
| --- | --- | --- | --- |
| **Stress response related** | Response to chitin | 49.66 | 0.00 |
|  | Response to stimulus | 43.96 | 0.00 |
|  | Response to carbohydrate stimulus | 43.59 | 0.00 |
|  | Response to chemical stimulus | 32.00 | 0.00 |
|  | Response to stress | 31.52 | 0.00 |
|  | Defense response | 31.15 | 0.00 |
|  | Response to organic substance | 29.59 | 0.00 |
|  | Immune response | 23.34 | 0.00 |
|  | Immune system process | 23.30 | 0.00 |
|  | Innate immune response | 19.47 | 0.00 |
|  | Response to other organism | 13.41 | 0.00 |
|  | Response to biotic stimulus | 13.32 | 0.00 |
|  | Defense response to fungus | 11.48 | 0.00 |
|  | Defense response to bacterium | 11.26 | 0.00 |
|  | Response to fungus | 11.09 | 0.00 |
|  | Response to bacterium | 10.60 | 0.00 |
|  | Response to wounding | 9.64 | 0.00 |
|  | Regulation of response to stress | 7.64 | 0.00 |
|  | Regulation of defense response | 7.49 | 0.00 |
|  | Response to abscisic acid stimulus | 6.85 | 0.00 |
|  | Indole glucosinolate metabolic process | 6.70 | 0.00 |
|  | Response to external stimulus | 6.62 | 0.00 |
| **Growth related** | Developmental growth | 0.00 | 2.18 |
|  | Cell morphogenesis | 0.00 | 2.06 |
|  | Cell growth | 0.00 | 2.02 |
|  | Growth | 0.00 | 1.60 |
|  | Developmental growth involved in morphogenesis | 0.00 | 2.66 |
|  | Unidimensional cell growth | 0.00 | 2.66 |
|  | Regulation of cell size | 0.00 | 1.96 |
|  | Regulation of cellular component size | 0.00 | 1.96 |
|  | Response to gibberellin stimulus | 0.00 | 2.96 |
|  | Cellular component morphogenesis | 0.00 | 1.89 |
|  | Cellular developmental process | 0.00 | 1.89 |
|  | Regulation of biological quality | 0.00 | 1.82 |
|  | Cellular component biogenesis | 0.00 | 1.68 |
|  | Ribosome biogenesis | 0.00 | 3.72 |
|  | Ribonucleoprotein complex biogenesis | 0.00 | 3.59 |

**Table S3. List of used primers.**

| **Gene** | **Primers** | **Sequence** | **Purpose** |
| --- | --- | --- | --- |
| ***PP2C.D1*** | Forward | GCTGC ATTGAAGGAA GCAGC | qRT-PCR |
|  | Reverse | TCA TGATGTTGAA TGCATCGGG |  |
| ***ALD1*** | Forward | GTGCAAGATCCTACCTTCCCGGC |  |
|  | Reverse | CGGTCCTTGGGGTCATAGCCAGA |  |
| ***FMO1*** | Forward | TCTTCTGCGTGCCGTAGTTTC |  |
|  | Reverse | CGCCATTTGACAAGAAGCATAG |  |
| ***SARD1*** | Forward | TCAAGGCGTTGTGGTTTGTG |  |
|  | Reverse | CGTCAACGACGGATAGTTTC |  |
| ***CBP60g*** | Forward | GATGACATGACCTCAAGCTG |  |
|  | Reverse | TTAACCTTACACCACCTGGC |  |
| ***SARD4*** | Forward | GCGAAACCAAGCTTGAGAAG |  |
|  | Reverse | TCCGGGTTTCAAGAACTCAC |  |
| ***ARF7*** | Forward | [TCAAGGTCACAGTGAGCAAGTCG](../../../../../about/blank) |  |
|  | Reverse | [TGTGGAGCATGCATATGAGCTTGG](../../../../../about/blank) |  |
| ***ARF19*** | Forward | [ACAGCTCGAAGATCCGCTAACC](../../../../../about/blank) |  |
|  | Reverse | [TGCACGCAGTTCACAAACTCTTC](../../../../../about/blank) |  |
| ***AT4G26410***  ***(Control gene)*** | Forward | GAGCTGAAGTGGCTTCCATGAC |  |
|  | Reverse | GGTCCGACATACCCATGATCC |  |
| ***ARF2*** | Forward | [CGTGAACAGGGAAGACCATTCCAG](../../../../../about/blank) |  |
|  | Reverse | [TTCCCTGCTTGTGAACCTTTGTG](../../../../../about/blank) |  |
| ***ARF3*** | Forward | [AGCCTGATATCCCTGTCTCTGAGG](../../../../../about/blank) |  |
|  | Reverse | [TTGACCTTGCAAGACCCTCTGG](../../../../../about/blank) |  |
| ***ARF10*** | Forward | [TGGCGACAACGTGAGAAAGACG](../../../../../about/blank) |  |
|  | Reverse | [AAGATGCTGAGCGGACCAGTCTTC](../../../../../about/blank) |  |
| ***ARF11*** | Forward | [CTATCTTCTCAATGGCCAGCTTCC](../../../../../about/blank) |  |
|  | Reverse | [ATGGCTCGTCCCATTGGATCTG](../../../../../about/blank) |  |
| ***ARF18*** | Forward | [TAAACACGCCACTGAATGCTTGCC](../../../../../about/blank) |  |
|  | Reverse | [AAATGCCTCCGTGGTTGTCCTCTG](../../../../../about/blank) |  |
| ***MPK3*** | Forward | CACC ATGAACACCG GCGGTGGCCA | Clone the CDS of the gene |
|  | Reverse | TTGCTGATATTCTGG |  |
| ***MPK6*** | Forward | CACC ATGGACGGTG GTTCAGGTCA | Clone the CDS of the gene without stop codon |
|  | Reverse | TTGCTGATATTCTGGATTGAA |  |
| ***PP2C.D1*** | Forward | ATGGTTAAACCCTGTTGGAGAATA | Clone the gene |
|  | Reverse | TCATGATGTTGAATGCATCGGG |  |
|  | Reverse | TGATGTTGAATGCATCGGG |  |
